# Bi-directional protein-protein interactions control liquid-liquid phase separation of PSD-95 and its interaction partners

**DOI:** 10.1101/2021.03.03.433781

**Authors:** Nikolaj Riis Christensen, Christian Parsbæk Pedersen, Vita Sereikaite, Jannik Nedergaard Pedersen, Maria Vistrup-Parry, Andreas Toft Sørensen, Daniel Otzen, Lise Arleth, Kaare Teilum, Kenneth L. Madsen, Kristian Strømgaard

**Author notes:** Corresponding authors &.

## Abstract

The organization of the postsynaptic density (PSD), a protein-dense semi-membraneless organelle, is mediated by numerous specific protein-protein interactions (PPIs) which constitute a functional post-synapse. Postsynaptic density protein 95 (PSD-95) interacts with a manifold of proteins, including the C-terminal of transmembrane AMPA receptor (AMAPR) regulatory proteins (TARPs). Here, we uncover the minimal essential peptide responsible for the stargazin (TARP-γ2) mediated liquid-liquid phase separation (LLPS) formation of PSD-95 and other key protein constituents of the PSD. Furthermore, we find that pharmacological inhibitors of PSD-95 can facilitate formation of LLPS. We found that in some cases LLPS formation is dependent on multivalent interactions while in other cases short peptides carrying a high charge are sufficient to promote LLPS in complex systems. This study offers a new perspective on PSD-95 interactions and their role in LLPS formation, while also considering the role of affinity over multivalency in LLPS systems.

## INTRODUCTION

Synaptic transmission is highly dependent on proper function and anchoring of ligand gated ion channels such as the α-amino-3-hydroxy-5-methyl-4-isoxazolepropionic acid receptors (AMPARs), which are responsible for the majority of fast excitatory transmission in the central nervous system (CNS). The postsynaptic density (PSD) contains ∼2000 proteins (Bayes et al., 2012; Bayes and Grant, 2009; Bayes et al., 2011; Distler et al., 2014; Grant, 2019; O’Rourke et al., 2012; Trinidad et al., 2008) and one of the most abundant proteins is postsynaptic density protein 95 (PSD-95, 724 residues [Human, Uniprot: P78352], molecular weight 80.5 kDa, with N-terminal palmitoylation 95 kDa) a master scaffold protein of the PSD. PSD-95 regulates the function of AMPARs indirectly through canonical and non-canonical PDZ-domain mediated interaction with several members of the TARP family, including Stargazin (Stg) also known as TARP-γ2, thereby anchoring the AMPAR/Stg receptor complex to the PSD membrane and further into the post synapse (Bissen et al., 2019; Zeng et al., 2019).

PSD-95 typically interacts with the C-terminus of a target protein through one of its three PDZ domains, and as is the case for most synaptic PDZ dependent interactions, the affinities are generally in the µM range (Christensen et al., 2019; Stiffler et al., 2007; Stiffler et al., 2006; Ye et al., 2018; Zhang et al., 2013). Nevertheless, several examples have emerged where PDZ binding is coupled to secondary binding sites, including the lipid membrane, which dramatically potentiates the affinity of the overall interaction (Erlendsson et al., 2019; Janezic et al., 2019; Zeng et al., 2019; Zeng et al., 2016a; Zeng et al., 2016b). Due to its role as a master scaffold protein in synaptic transmission, PSD-95 has been suggested as a promising drug target for treatment of ischemic stroke and chronic pain amongst others. Currently, there are several lead candidates targeting PSD-95 in both pre-clinical and clinical development (Christensen et al., 2019), covering both small molecules (Florio et al., 2009; Hu et al., 2013; Lee et al., 2015; Wu et al., 2014) and in particular peptide derived compounds (Bach et al., 2009; Bach et al., 2012; Bach et al., 2011; Long et al., 2003; Nissen et al., 2015; Piserchio et al., 2004; Sainlos et al., 2011; Aarts et al., 2002), all of which target the PDZ domains of PSD-95. These molecules feature both monovalent and multivalent interactions with PSD-95, and their affinities range from µM (Florio et al., 2009; Piserchio et al., 2004; Aarts et al., 2002) to low nM (Bach et al., 2009; Bach et al., 2012; Nissen et al., 2015).

Recently, liquid-liquid phase separation (LLPS) and the formation of membraneless organelles have emerged as a common feature of protein assembly in many branches of cellular biology (Alberti et al., 2019; Banani et al., 2017). It was recently shown that PSD-95 can undergo LLPS in different ways, both in complex with synaptic Ras GTPase-activating protein (SynGAP) and in complex with four additional proteins, namely homer protein homolog 3 (Homer3), SH3, multiple ankyrin repeat domains 3 (Shank3) and guanylate kinase-associated protein (GKAP). In addition, PSD-95 also undertake LLPS in complex with the Stg C-terminus and the C-termini of other members of the TARP family as well as the C-terminal of N-methyl-D-aspartate receptors (NMDAR) (Tao et al., 2019; Zeng et al., 2018; Zeng et al., 2019; Zeng et al., 2016a). The LLPS of the key PSD components suggest that the formation of a postsynaptic condensate can govern key aspects of synaptic transmission (Zeng et al., 2019).

In this study we investigated the behavior of PSD-95 in solution and in complex with multivalent ligands derived from the Stargazin C-terminal region. We show how PSD-95 behaves as a monomer in solution and that PSD-95 can undergo LLPS in absence and presence of ligands and key components of the PSD. We then show how PSD-95 can act as a bi-directional modulator of LLPS formation and confirm a secondary charge dense binding site in Stargazin for PSD-95. Finally, we tested the ability of known, charge-dense, pharmacologically relevant inhibitors of PSD-95 to induce LLPS. Taken together our findings suggest novel mechanisms for ligand-induced PSD-95 LLPS formation, defined by highly charged peptides and/or multivalent interactions.

## RESULTS

### PSD-95 organizes as a monomer with compact tertiary structure in solution

Structural studies of PSD-95 have previously shown that PSD-95 can organize as either an elongated protein arranged perpendicular to the synaptic membrane (Chen et al., 2011) or in a more compact fashion in negative stain EM (Fomina et al., 2011). High-resolution structures of all the individual structured domains of PSD-95 have been obtained (Bach et al., 2012; Long et al., 2003; McGee et al., 2001; Rodzli et al., 2020; Sainlos et al., 2011; Tavares et al., 2001; Zeng et al., 2016b; Zhu et al., 2017), while the complex interplay and dynamics in solution between the structured domains and flexible linkers remain to be explored (Figure 1A). To bridge this structural information, we investigated the solution structure of full-length PSD-95 using small angle X-ray scattering (SAXS).

**Figure 1.**
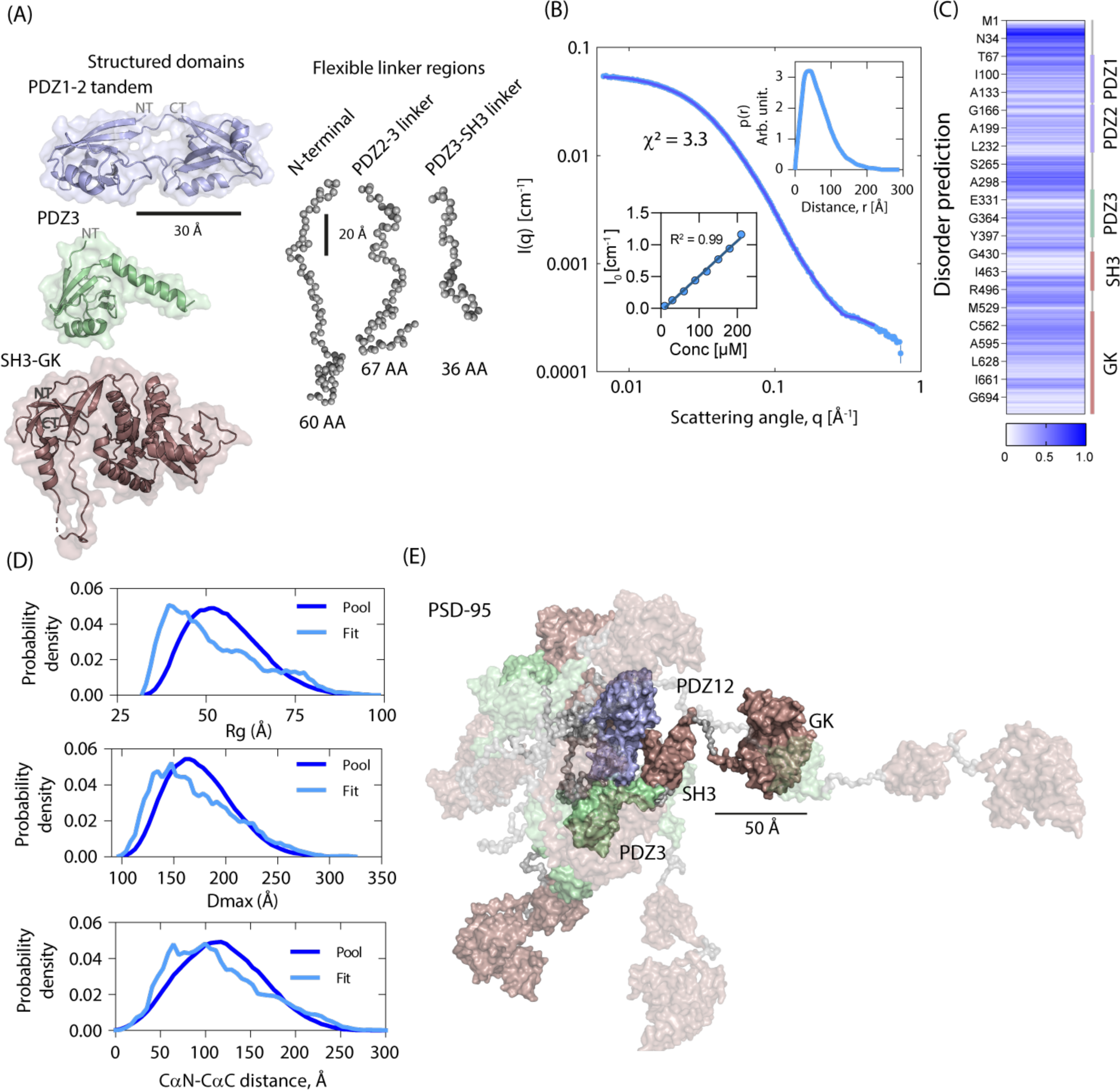
PSD-95 arranges in a compact state in solution and can undergo pH dependent LLPS. (A) Structured domain and unstructured regions of PSD-95 (PDB: 3GSL, 5JXB, 1KJW). (B) SAXS data for PSD-95 at 60 µM, with I(0) vs concentration dependency and pair-distance distribution, *p(r)*, (GenApp.Rocks) as inserts. Fit and χ^2^ indicates fit of EOM modeling to the data. (C) Prediction of PSD-95 disorderedness (IUpred2A) (Dosztanyi, 2018; Dosztanyi et al., 2005; Meszaros et al., 2018; Meszaros et al., 2009) with domains indicated to the right. (D) EOM obtained parameters for ensemble, with R_g_ distribution (top panel), D_max_ distribution (middle panel) and CαN-CαC distance distribution (bottom panel) suggest partial domain flexibility and a relatively compact structure of PSD-95 in solution. (E) Proposed EOM ensemble for PSD-95 with shading according to model ensemble percentage (8.1/8.1/8.1/8.1/8.1/17.2/17.1/25.3%). The disordered N-terminals are removed for clarity.

We found that full length PSD-95 is stable in solution, with a mean radius of gyration (R_g_) = 59 Å ± 4 Å and a maximal distance (D_max_) = 245 Å ± 23 Å, and a M_w,calc_ = 88 kDa ± 8 kDa which corresponds well with a monomer of PSD-95 (M_Wteo_ = 80.8 kDa), across the 30-210 µM concentration span (Figure 1B insert, and Figure S1A). A Kratky plot (Kikhney and Svergun, 2015) of the scattering data (Figure S1B) suggested that PSD-95 is a flexible protein, as would be expected, since PSD-95 comprises a series of protein domains connected by flexible linkers and a disordered N-terminus (Figure 1A-B and Figure S1B). Hence, we used an ensemble optimization method (EOM) (Bernado et al., 2007; Tria et al., 2015) to fit an ensemble of PSD-95 conformations (Figure 1D-E). This provides a better representation of the flexible structures than a single state model. To obtain a solution structure of full-length PSD-95, we generated a series of models using known structural domains of PSD-95 (PDZ1-2, PDB: 3GSL; PDZ3, PDB: 5JXB; SH3-GK, PDB: 1KJW), connected via fully-flexible linkers. Using these input structures, we generated 10.000 conformations using EOM (Bernado et al., 2007; Tria et al., 2015) (Ranch), and used the genetic algorithm (GAJOE) to fit the data recorded at a protein concentration of 60 µM. This yielded a subset of structures representing the final structural ensemble (Figure 1D and Figure 1E) (Bernado et al., 2007; Tria et al., 2015). The final ensemble had an R_g, mean_ = 52.8 Å and a D_max,mean_ = 173.4 Å. The final structural ensemble points towards that PSD-95 on average adopt more compact conformations in that proposed by the generating ensemble. We note however that it was necessary to include also extended structures to fit the obtained SAXS data and around 16% of the structural ensemble had conformations with a D_max_ above 200 Å (Figure 1D). This behavior is in overall agreement with previously published structural data suggesting that PSD-95 both forms extended and compact conformations (Fomina et al., 2011).

### Multivalent Stg peptides induce LLPS condensate formation when mixed with PSD-95

At the PSD, PSD-95 is always bound to other proteins such as the AMPAR auxiliary subunit Stargazin (Stg), the GluN2B subunit of the NMDAR, adhesion proteins, such as Neuroligns or other scaffold proteins such as GKAP or SynGAP. Several of the interactions with PSD-95 are multivalent, often due to oligomeric protein assemblies. An example is Stg, which forms a complex with AMPAR, where the Stg:AMPAR stoichiometry can be from 1:1 to 4:1 (Twomey et al., 2016; Zhao et al., 2016). The Stg C-terminal has previously been shown to interact with all the PDZ domains of PSD-95, with a preference for the PDZ1-2 tandem over PDZ3 (Pedersen et al., 2017; Sainlos et al., 2011). To mimic the above mentioned difference in oligomeric states for the Stg:AMPAR complex, we designed peptides which in solution organize as monomers, dimers or trimers, thereby varying the number of available PDZ binding motifs (PBMs) targeting PSD-95 between 1 and 3 (Figure 2A). The dimeric variant was designed using the General control protein GCN4 (GCN4) leucine zipper motif, previously found to form a homo-dimeric parallel helical leucine zipper in solution (Gonzalez et al., 1996), fused to a hexapeptide corresponding to the 6 C-terminal residues of Stg (RRTTPV), through a short flexible linker, yielding a dimeric Stg C6 variant (dim-Stg, Figure 2A). To disrupt the helical GCN4p1 motif, we inserted two prolines (mono-Stg), yielding a monomeric conformation of the Stg C-terminal (Hanes et al., 1998; Leder et al., 1995). To conversely increase the number of PBMs, we also made a trimeric GCN4p1 variant (tri-Stg, Figure 2C) (Harbury et al., 1994). As expected, we found that both dim-Stg and tri-Stg displayed a high degree of helicity in solution, wherein the tri-Stg has an even higher degree of helicity than that of dim-Stg, presumably due to more cooperative folding, combined with a lower elution volume for tri-Stg than dim-Stg in size exclusion chromatography, suggesting a larger hydrodynamic radius, while the monomer, mono-Stg, displayed a random coil like structure, and an elution volume similar to dim-Stg (Figure S2A-B). Using competitive fluorescence polarization (FP) binding to full length PSD-95 (Zeng et al., 2016b) we found that mono-Stg, dim-Stg and tri-Stg had apparent affinities of K_i_ = 7984 nM (SEM: [7146;8910] nM, n=6), K_i_ = 237 nM (SEM: [195;289] nM, n=6) and K_i_ = 98 nM (SEM: [81;119] nM, n=6), respectively (Figure 2B). This demonstrates an affinity gain of 33- and 81-fold for dim-Stg and tri-Stg, respectively over mono-Stg, which is comparable to the 25-fold affinity gain seen for prior work on bivalent Stg peptides (Sainlos et al., 2011).

**Figure 2.**
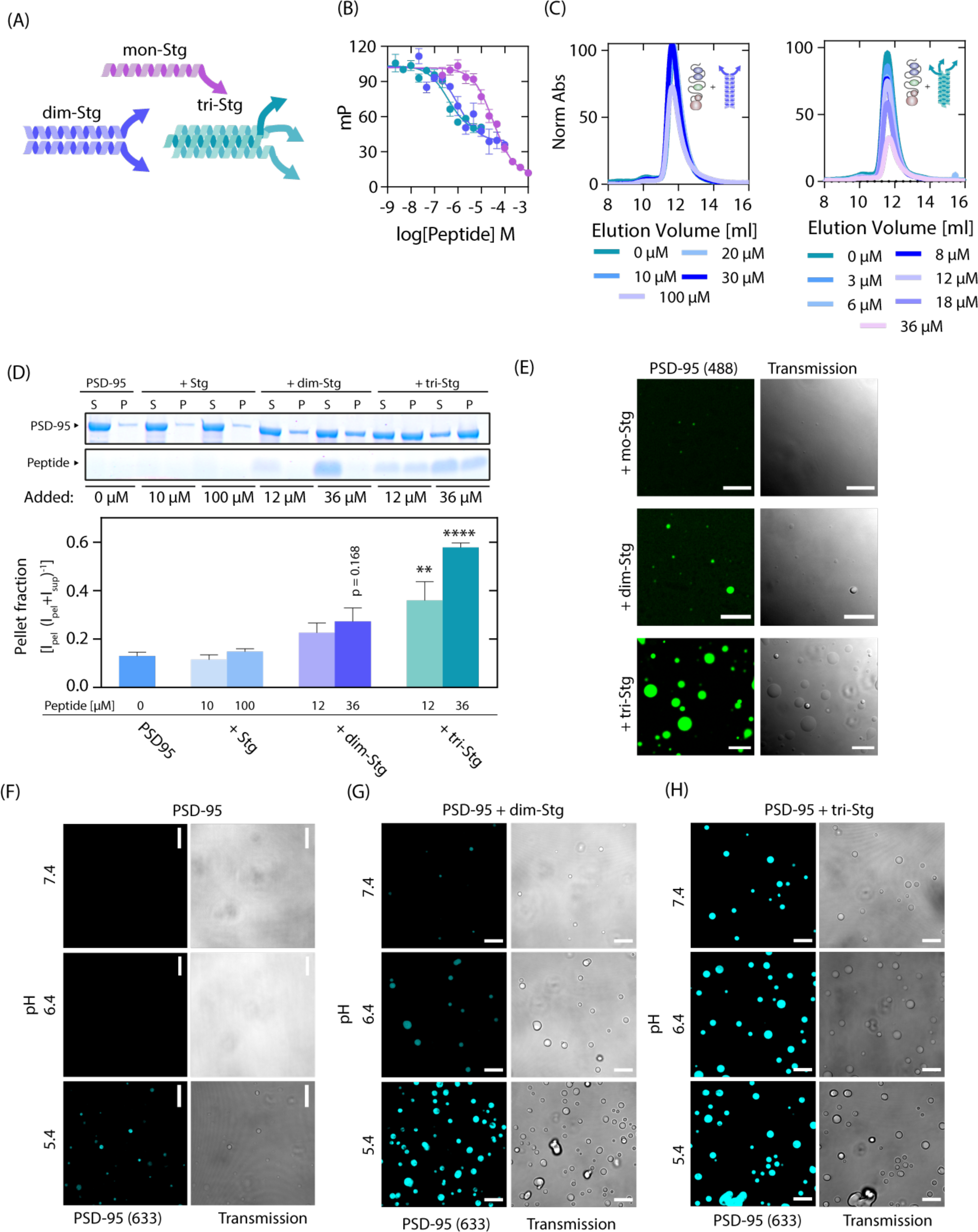
Multivalent PSD-95-peptide interactions can induce concentration and pH dependent LLPS. (A) Illustration of tested monomeric, dimeric or trimeric peptides. (B) Fluorescence polarization competition with full length PSD-95 shows 33-fold and 81-fold increased affinities for dim-Stg (K_i_ = 237 nM SEM [195;289] nM, n = 6) and tri-Stg (K_i_ = 98 nM, SEM [81;119] nM, n = 6) over mono-Stg (K_i_ = 7984 nM, SEM [7146;8910] nM, n=6) respectively. (C) Size exclusion chromatography elution profile of PSD-95 (10 µM) incubated with increasing amounts of dim-Stg or tri-Stg. Traces were extracted as absorbance at 280 nm and normalized to the elution of PSD-95 in absence of peptide. (D) SDS-PAGE sedimentation assay with full length PSD-95 incubated with Stg C11, dim-Stg or tri-Stg indicates formation of liquid-liquid phase separation (LLPS) condensates for dim-Stg and tri-Stg, but not Stg-C11. (E) LLPS formation was verified for dim-Stg and tri-Stg by confocal microscopy using fluorescently labeled PSD-95 incubated with 36 µM of mono-Stg, dim-Stg and tri-Stg. Error bars are shown as SEM of (B) n=6 or (D) n=3. Statistics was done using one-way ANOVA with Dunnett post-test. *, p<0.05; **, p< 0.01; ***, p<0.001; **** p<0.0001. (F-H) pH dependent LLPS formation was seen for PSD-95, in absence or presence of dim-Stg (G) or tri-Stg (H) using confocal microscopy visualized using fluorescently labeled Alexa-6-PSD-95.

Size exclusion chromatography (SEC) and SEC multiangle light scattering (SEC-MALS) demonstrated that incubation with either of the peptides did not change the elution volume or molecular weight of PSD-95 (Figure 2C and Figure S2C-F). To our surprise, however, we found that an increase in dim-Stg and tri-Stg concentration with a fixed PSD-95 concentration, caused a reduction in the total amount of PSD-95/peptide complex eluting from the column (Figure 2C). This was validated using SEC-MALS, also substantiating that no oligomeric PSD-95 eluted from the column (Figure S2C-F). Using Flow induced dispersion analysis (FIDA) (Pedersen et al., 2019) we found that the hydrodynamic radius (RH) was seemingly larger for the Stg-bound complexes than it was for PSD-95 in absence of the Stg peptides, however the RH increase was only significant at 36 µM tri-Stg (*, p=0.011, one-way ANOVA) (Figure S2G). Data spikes in the FIDA data also indicated a presence of aggregates which we did not see in SEC or SEC-MALS, suggesting that the oligomers formed were too large to enter the SEC columns (Figure S1H). We investigated this phenomenon further using a SDS page protein sedimentation assay (Wu et al., 2019; Zeng et al., 2018; Zeng et al., 2019; Zeng et al., 2016a). We found that both dim-Stg and tri-Stg, but not monomeric Stg, induced a cloudy phase which could be pelleted upon centrifugation (Figure 2D). The pellet induction was significant for tri-Stg at 12 µM (**, p<0.01, one-way ANOVA, Dunnett post-test) and 36 µM (****, p<0.0001 one-way ANOVA, Dunnett post-test) (Figure 2D). To evaluate if the Stg C-terminal peptides induced a LLPS transition, we performed fluorescence confocal microscopy of Alexa488-labeled PSD-95 and unlabeled peptides. Indeed, we found that mixing dim-Stg and tri-Stg (at 36 µM) with PSD-95 (3 µM) induced LLPS droplets (Figure 2E). The formation of LLPS droplets suggested, that multivalent peptides containing multiple copies of a PBM targeting PSD-95 PDZ domains such as the Stg peptides, are sufficient to induce LLPS when mixed with PSD-95.

Synaptic function is highly dependent on the pH homeostasis on both the extra- and intracellular side of the postsynaptic membrane. The effect of pH alterations on the protein complex formation in the PSD is currently unknown. We therefore moved on to investigate the effects of pH on the LLPS assembly of PSD-95 and the Stg peptides. Upon acidification of the buffer to pH 5.4, we observed formation of a hazy precipitate in the sample tube suggesting sample precipitation (not shown). Using fluorescence confocal microscopy, we found that the precipitate, probed with Alexa633-labeled PSD-95 behaved as dynamic droplets (Figure 2F), suggesting that PSD-95 can spontaneously undergo LLPS formation in a pH dependent manner, also strengthening the conception that PSD-95 can self-organize in larger oligomers in a ligand independent manner. We saw that the formed droplets were dynamic in size, and fluorescence recovery after photo bleaching (FRAP) experiments consistently demonstrated partial recovery (Figure S2I-K).

A recent NMR based study showed that PDZ1-2 can self-associate, and indeed we also saw that at high concentrations (1.45 mM) that most amide proton resonances in the PDZ1-2 ^1^H-^15^N-HSQC experienced severe line broadening, to an extent where numerous peaks disappeared (Figure S3A). The line broadening was most likely caused by dynamic processes between individual PDZ1-2 proteins in the intermediate NMR time scale, since these were concentration dependent (Figure S3A), and the peaks reappeared at a lower concentration. We mapped the concentration induced intensity changes (Figure S3B) and found that the charged and non-polar residues for which the intensity is more than 30% reduced or increased, accounted for 82% of the residues, compared to 18% for the polar non-charged residues. Once mapped onto the structure of PDZ1-2 (PDB 3GSL; Figure S3C), we found that all the residues, with an intensity changes of more than 30% in either direction, were surface exposed and most of the residues were located on the opposite side relative to the binding pocket of both PDZ domains, suggesting that these might be involved in PDZ1-2/PDZ1-2 protein interactions, as have also been suggested from earlier structural work done on the PDZ1-2 tandem (Rodzli et al., 2020; Sainlos et al., 2011). This may suggest that the PDZ1-2 display several low affinity interactions that may facilitate PSD-95 oligomerization.

We next tested whether the ability of dim-Stg and tri-Stg and pH works in an additive manner to induce LLPS (Figure 2G-H). We found that upon acidification of the buffer there was a pronounced positive effect on LLPS formation for both dim-Stg and tri-Stg, this was also shown by SDS-PAGE sedimentation, where it was evident that tri-Stg enhanced the protein content in the pellet both at pH 6.4 and 5.4 (Figure S2L).

Taken together the solution structure of PSD-95 combined with the observation that PSD-95 can undergo LLPS at acidic pH suggests the presence of weak intra protein interactions both within a single PSD-95 protein and between individual PSD-95 proteins, mediated in part by the PDZ1-2 tandem of PSD-95.

### The PDZ1-2 tandem of PSD-95 is sufficient to cause LLPS when mixed with Stg peptides

Since PSD-95 is a multi-domain protein, we wanted to evaluate if the PDZ1-2 tandem of PSD-95 could provide a protein scaffold of sufficient valency to promote LLPS. We found that dim-Stg and tri-Stg could induce LLPS of Alexa488-labeled PDZ1-2 alone, as seen from fluorescence confocal microscopy using fluorescence microscopy (Figure 3A and 3B) and SDS-PAGE sedimentation (Figure S2A), similar to the findings for full-length PSD-95. To investigate the residues involved in LLPS interaction network with dim-Stg and tri-Stg, we recorded ^1^H-^15^N-HSQC NMR experiments of PSD-95 in complex with the two peptides (Figure 3C and 3D). The majority of resonances of the two samples experienced severe line-broadening compared to the absence of the peptides, which is likely caused by the dynamic properties of the interaction networks in the phase separated droplets. We used SEC-MALS to estimate the binding valency, and found that both the elution volume and the molecular weight of the eluting complex are consistent with a 1:1 stoichiometry of the interaction between PDZ1-2 and the Stg peptides at a concentration below the critical LLPS concentration (Figure 3E and 3F). Taken together these data support that the tandem PDZ1-2 protein combined with dim-Stg or tri-Stg is sufficient for LLPS formation and suggest that the LLPS core of PSD-95 is the PDZ1-2 tandem.

**Figure 3.**
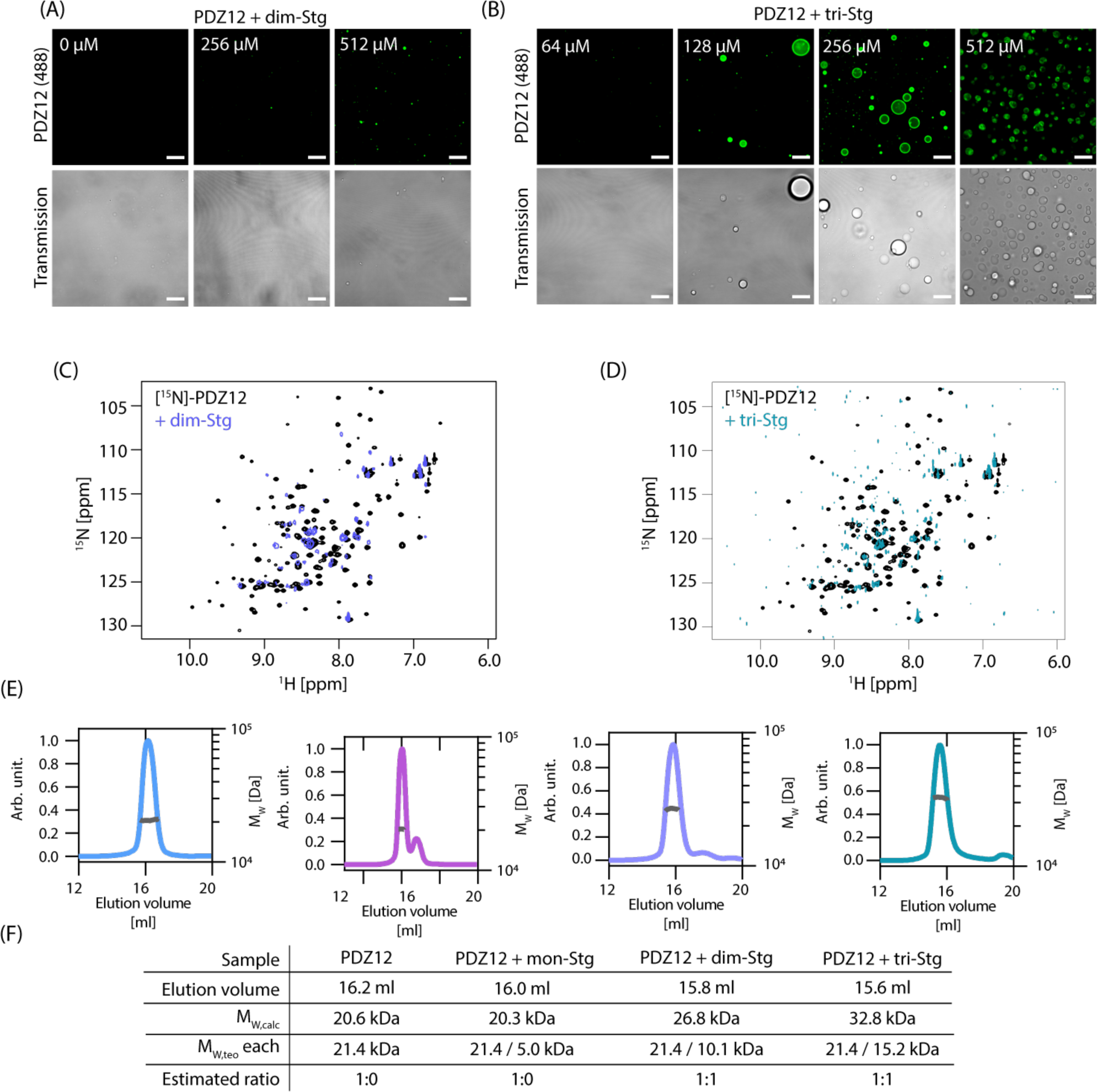
Simple stoichiometric binding of multivalent interaction partners leads to LLPS formation of the PDZ12 tandem from PSD-95. (A) Concentration dependency of LLPS induction for PDZ1-2 incubated with dim-Stg show only minor LLPS induction. (B) Concentration dependency of LLPS induction for PDZ1-2 incubated with tri-Stg show LLPS induction at peptide:protein ratios above 1:1 and suggests a biphasic droplet formation. (C-D) [^1^H]-[^15^N]-HSQC spectra overlay of 100 μM [^15^N]-labelled PSD-95 PDZ12 (black) with 512 μM dim-Stg (C) and 512 µM tri-Stg (D), show severe line-broadening likely caused by the dynamic property of the interaction network in the phase separated droplets. (E) SEC-MALS elution profiles and molecular weight calculation of 200 µM PDZ1-2 (blue) incubated with 600 µM mono-Stg (purple), dim-Stg (blue) or tri-Stg (teal). (F) Data table of fitted data from (E), indicating 1:1 complexes between PDZ1-2 dim-Stg or tri-Stg.Fitting was done using ASTRA and data plotting was done using GraphPad Prism 8.3.

### PSD-95 serves as a reversible, negative modulator of condensate formation governed by multivalent PDZ interactions

Based on earlier observations of the five major synaptic scaffold proteins (PSD-95, Homer3, Shank3, GKAP and SynGAP, *see Methods*) (Zeng et al., 2018; Zeng et al., 2016a), and the formation of LLPS droplets upon mixing (Zeng et al., 2018; Zeng et al., 2019; Zeng et al., 2016a), we expressed and purified these five major synaptic scaffold proteins. We found that, in absence of PSD-95, the Homer3, Shank3, GKAP and SynGAP (H-S-G-S) complex, condensed into LLPS droplets (3 µM each) (Figure 4A). Surprisingly, upon incubation with increasing amounts of PSD-95, this phase separation was significantly reduced in presence of PSD-95 (3 µM) (*** for Homer3, p<0.001; ** for Shank3, p<0.01; * for GKAP p=0.0113; ** for SynGAP, p<0.01; two-way ANOVA Dunnett post-test) and at 10 µM of PSD-95 (** for Homer3, p<0.01; ** for Shank3, p<0.01; * for GKAP p=0.013; ** for SynGAP, p<0.01; two-way ANOVA Dunnett post-test) (Figure 4B). This was confirmed using confocal microscopy, where we observed LLPS droplets formed by 3 µM of H-S-G-S in the absence of PSD-95 (Figure 4C) while only minor droplets were seen in the presence of PSD-95 (1 µM) (Figure 4D). This effect was also evident upon addition of PSD-95 (8 µM) to pre-existing H-S-G-S condensates (3 µM each), probed with Alexa647-labeled Shank3 (0.3 µM) and Alexa488-labeled PSD-95 (0.8 µM) (Figure 4E). When PSD-95 associated to the droplets, there was a slow incorporation of PSD-95 into the droplets from the periphery that gradually reduced Shank3 intensity (Figure 4F), which was quantified to ∼30% reduction in Shank3 intensity (Figure 4G). No effect was seen upon addition of PBS (Figure S4A-B) indicating a PSD-95 dependent disassembly of the H-S-G-S condensate.

**Figure 4.**
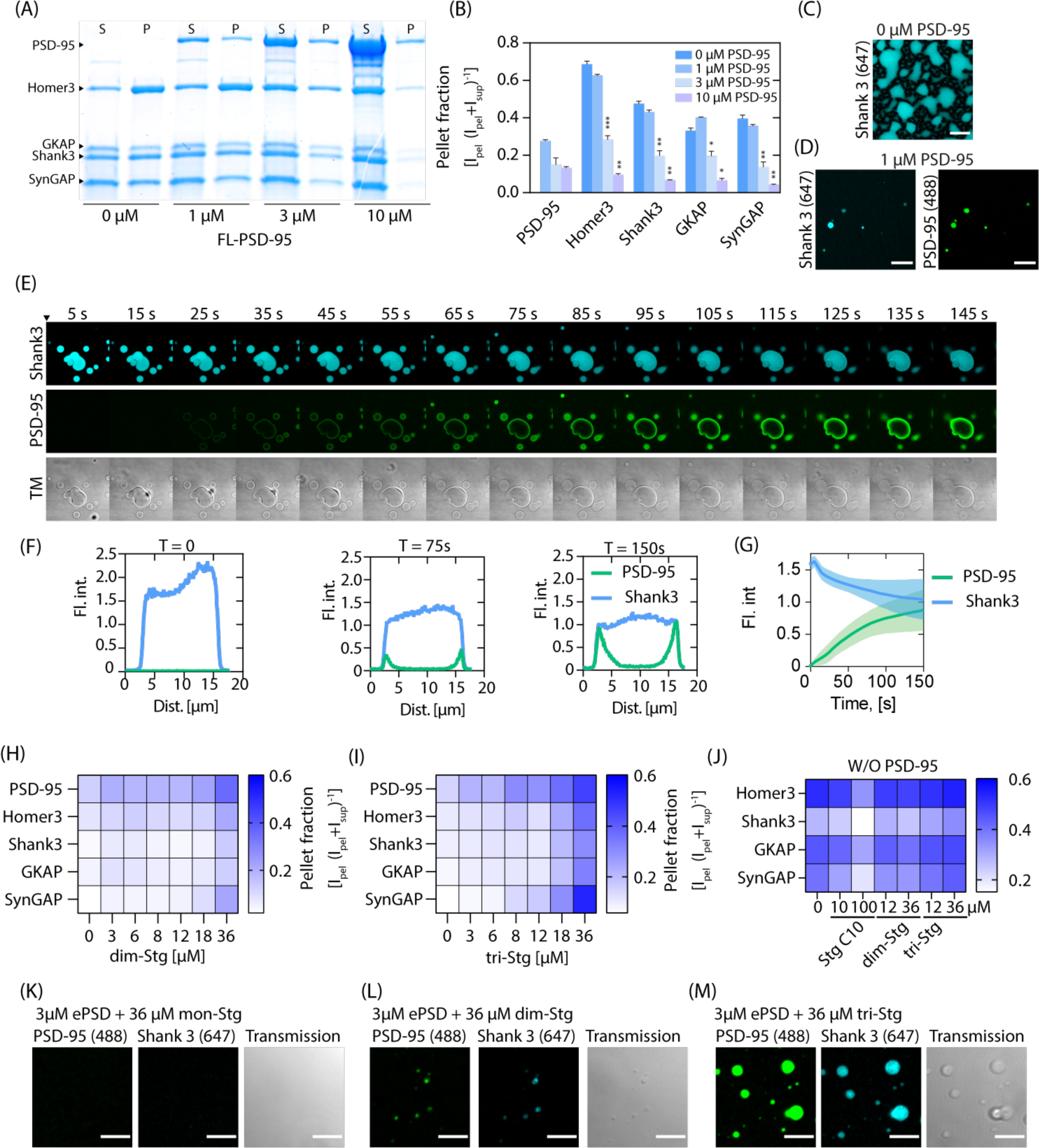
ePSD condensate can be modulated through multivalent PSD-95 PDZ interactions. (A) Representative SDS-PAGE gel of sedimentation assay with 3 µM Homer3, Shank3, GKAP and SynGAP (H-S-G-S) incubated with increasing amounts of PSD-95. (B) Quantification of the gels shown in (A) shows reduction of condensate formation as a function of PSD-95 addition. (C) Validation of H-S-G-S LLPS condensate formation using confocal microscopy, with fluorescently labelled Shank3, scalebars 10 µm. (D) Confocal microscopy of 5x ePSD condensate with fluorescently labelled PSD-95 and Shank3, scalebar 10 µm. (E) Confocal microscopy time series of H-S-G-S condensate upon addition of PSD-95 indicates slow absorption of PSD-95 into existing droplets. (F) Line intensity profile of PSD-95 (Green) and Shank3 (blue) at indicated timepoints, which show a time dependent reduction in Shank3 signal. (G) Quantification of mean droplet intensity of Shank3 (blue) and PSD-95 (green) after addition of PSD-95. Error band is the SEM of three independent chambers. (H-I) Heatmap of SDS-PAGE sedimentation quantification of 5xePSD (3 µM H-S-G-S, 10 µM PSD-95) incubated with increasing amounts of dim-Stg (H) or tri-Stg (I). (J) Heatmap representation of SDS-PAGE sedimentation quantification of H-S-G-S condensate (3 µM H-S-G-S) incubated with Stg C10, dim-Stg or tri-Stg. (K-M) confocal microscopy confirmation of LLPS formation upon addition of dim-Stg and tri-Stg (36 µM) to ePSD (3 µM). Error bars are shown as SEM of n=3, scalebars 5 µm

We next asked how the di- and trimeric Stg peptides would affect the destabilized condensate of all five major PSD proteins (ePSD). Indeed, condensate formation was facilitated with increasing concentration of dim-Stg and, in particular, tri-Stg (Figure 4H-I and Figure S5A-D), but not mono-Stg (Figure S4C and S5E-F), suggesting that the peptide valency and concentration is critical for ePSD condensate stabilization. The induction of LLPS in presence of dim-Stg and tri-Stg was validated by confocal microscopy, which suggested that tri-Stg was much more efficient than mono-Stg and dim-Stg at stabilizing LLPS in the ePSD (Figure 4K-M). While dim-Stg and tri-Stg could be used to induce LLPS in the ePSD complex, the H-S-G-S condensates were unaffected by dim-Stg and tri-Stg, while high concentrations of mono-Stg seemed to reduce pelleting of the H-S-G-S condensate (Figure 4J and Figure S5G-H).

These data demonstrate a critical role of PSD-95 in the modulation of the ePSD condensates, which in turn is governed by specific multivalent PDZ domain interactions.

### Stg contains multiple PSD-95 binding sites

It was recently shown that the Stg C-terminus in its full length can induce LLPS when mixed with PSD-95 alone or in combination with Homer3, Shank3, GKAP and SynGAP. This effect is inhibited by S-to-D mutations in an S/R rich region (S221-S253, Uniprot: Q9Y698) positioned, upstream of the PBM (T321-V323), in the membrane proximal region (Feng et al., 2019b; Zeng et al., 2019). The Stg C-terminus (Figure S6A) was shown to interact with PDZ2 through the PBM and non-canonically with PDZ1 via the S/R rich region probably interacting with E/D residues positioned opposite to the canonical PDZ1 binding pocket in PDZ1 (Zeng et al., 2019). To characterize the non-canonical PDZ interaction of PSD-95 with the Stg C-terminus, we designed and prepared a celluSPOT array (Winkler et al., 2009; Wu and Li, 2009) of the entire Stg C-terminus consisting of 101 15-mer peptides (Figure 5A), consecutively shifted one residue towards the C-terminus. The celluSPOT approach involves coupling of the C-terminal carboxylic acid to the cellulose membrane, thereby blocking canonical PDZ interactions, allowing us to probe non-canonical PDZ interactions of the Stg C-terminus with PSD-95. In addition, we included 39 peptides, carrying the S-to-D mutations in the S/R rich region described earlier (Sumioka et al., 2010; Tomita et al., 2005), to address putative modulation by phosphorylation.

**Figure 5.**
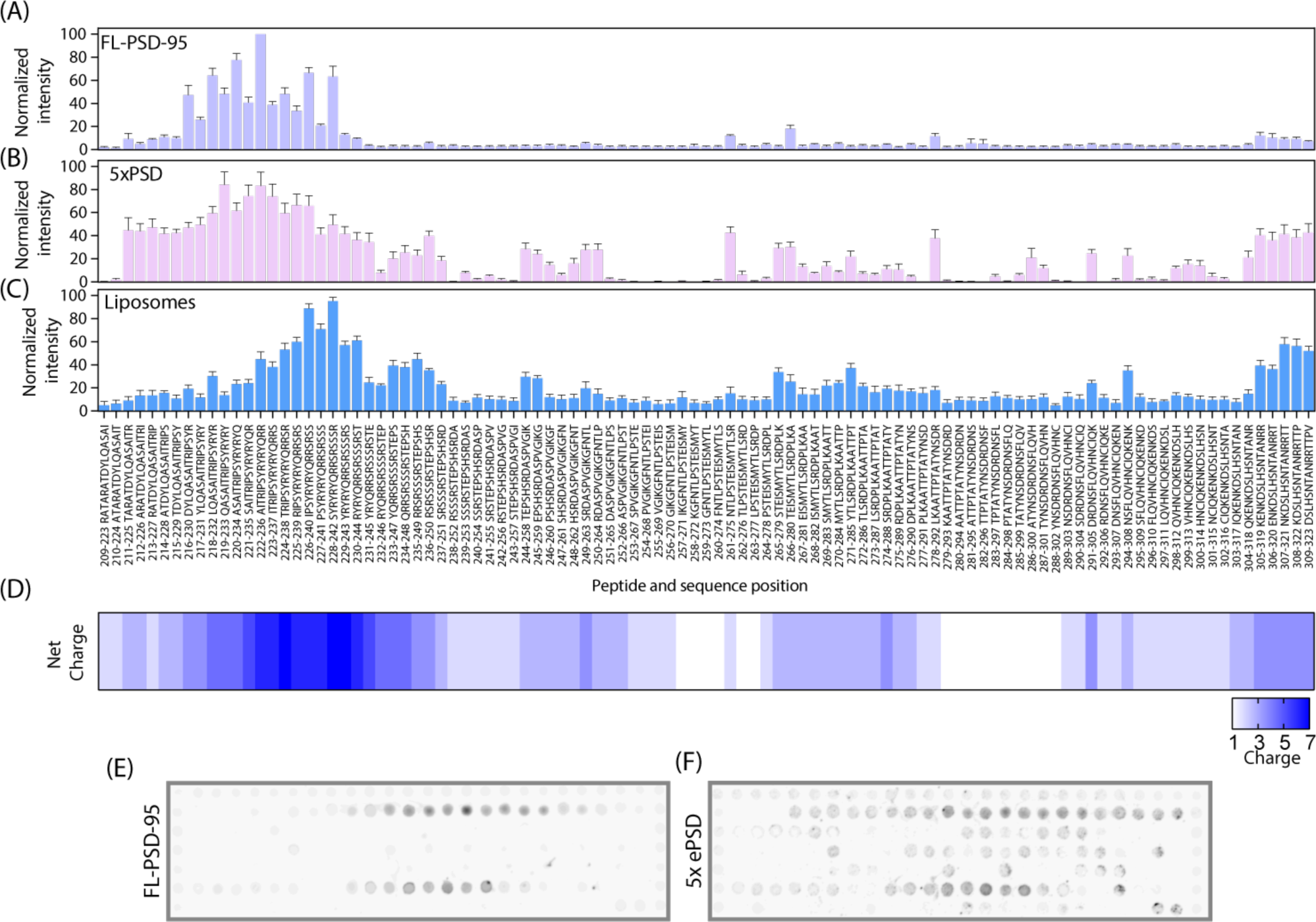
PSD-95 and ePSD condensate relies on a binding site outside the PDZ motif. (A-C) Quantification of CelluSPOT arrays of Stg C-terminal peptides (16-mers) when incubated with PSD-95 (A), ePSD (B) or vesicles (C). Primary sequence of peptides is indicated below each bar. Error bars are shown as SEM of n=8 for PSD-95 binding, n = 6 for ePSD and n = 8 for vesicles. (D) Heatmap of peptide net charge. (E-F) Representative celluSPOT array membranes of Stg C-terminal as 16-mer peptides, with a single residue frame shift per peptide, when incubated with PSD-95 (1 µM, (E)) or ePSD (3 µM H-S-G-S and 1 µM PSD-95, (F)).

We found that the intensities in the membrane proximal region of Stg (A214-E245), which overlaps partially with the charged S/R rich region (Figure 5A), were selectively enhanced upon addition of PSD-95 (1 µM total, 0.9 µM unlabeled PSD-95 and 0.1 µM Alexa633-labled PSD-95), suggesting a secondary binding site for PSD-95 in the A214-E245 region (Figure 5A and Figure S6B). Our peptide array further suggested that the S-to-D variants (Figure S6C) slightly reduced the binding to isolated PSD-95 in the membrane proximal region, confining the binding region from residues T211-S253 to T215-D241 (Figure S6D). To validate the binding of the S/R rich region of Stg to PSD-95, we then performed FP on a fluorescently labeled 15-mer peptide (TAMRA-G-A222ITRIPSYRYRYQRR_236_) representing the core binding region and found that PSD-95 indeed bind to Stg_A222-R236_ (K_d_ = 14.9 ± 3.0 µM) (Figure S6E). Interestingly, the S/R rich region overlaps with binding site of Arc (Zhang et al., 2015), which also show binding to Stg_A222-R236_ (K_d_ = 8.37 ± 1.25 µM) (Figure S7A-B).

Next, adding the remaining components of the ePSD (1 µM of H-S-G-S and 0.9 µM unlabeled PSD-95 and 0.1 µM Alexa633-labled PSD-95) to the Stg array caused a potentiation and broadening of the PSD-95 signal now covering the T211-S253 region as well as at the extreme C-terminus (Q304-V323) of Stg (Figure 5B). All ePSD proteins were also tested individually for binding to the Stg C-terminal peptides, and this resulted in a measurable intensity in the same membrane proximal region for all proteins (Figure S8A), suggesting a low specificity interaction, which was supported by FP binding data (Figure S8B). Moreover, the negative effect of the S-to-D mutations on PSD-95 binding was maintained in presence of the ePSD complex, in particular in the R225-A252 region (Figure S8C).

The Stg C-terminus has previously been shown to interact also with lipid membranes in an S-to-D dependent manner. We therefore tested liposome binding to the array and found that the areas Q219-D251, partially overlapping with the protein binding region, as well as the C-terminal region Q304-V323 showed liposome binding (Figure 5C), in accordance with a previously reported lipid binding site of the Stg C-terminal (Sumioka et al., 2010). This binding largely reflected the charge distribution (Figure S8E) and accordingly it was potently modulated by S-to-D mutations (Figure S6D and S8C-D).

Taken together, the peptide array, binding experiments suggest that the secondary PSD-95 binding site in Stg C-terminus is confined to the region A214-E245, which largely overlaps with the lipid binding site covering the region Q219-D251 (Sumioka et al., 2010) and is consistent with the suggestion that the S/R rich region is also involved in the binding of PSD-95 in a way which enhances affinity (Zeng et al., 2019).

### Stg_A222-R236_ can induce LLPS

As the Stg C-terminal was recently shown to induce LLPS in an R and S-to-D dependent manner, we wondered whether the 15-mer Stg_A222-R236_ peptide, which is part of the S/R rich region, was sufficient to induce LLPS. Indeed, we found that the Stg_A222-R236_ peptide (100 µM) induced LLPS for the ePSD (3 µM) (Figure 6A-B, and 6E). However, Stg_A222-R236_ (100 µM) did not induce LLPS for PSD-95 alone (3 µM) at pH 7.4 but LLPS was observed at the slightly more acidic pH 5.4 (Figure 6D and Figure S9B), similar to what was observed for PSD-95 in absence of ligands. As demonstrated above, the addition of PSD-95 to existing H-S-G-S droplets resulted in a reduction in Shank3 intensity, while PSD-95 was slowly taken up into the droplets from the periphery of preexisting droplets (Figure 4E). When adding Stg_A222-R236_ to H-S-G-S droplets, which had first been subjected to addition of PSD-95, we found that Stg_A222-R236_ was taken up rapidly by existing droplets (Figure 6E) with a flat gradient across the droplets (Figure 6G-I), as opposed to the peripheral localization seen for PSD-95 (Figure 4E-G). Further, this peripheral localization for PSD-95 was compromised in presence of Stg_A222-R236_ droplets, suggesting alterations in the dynamic protein network (Figure 6G-I).

**Figure 6.**
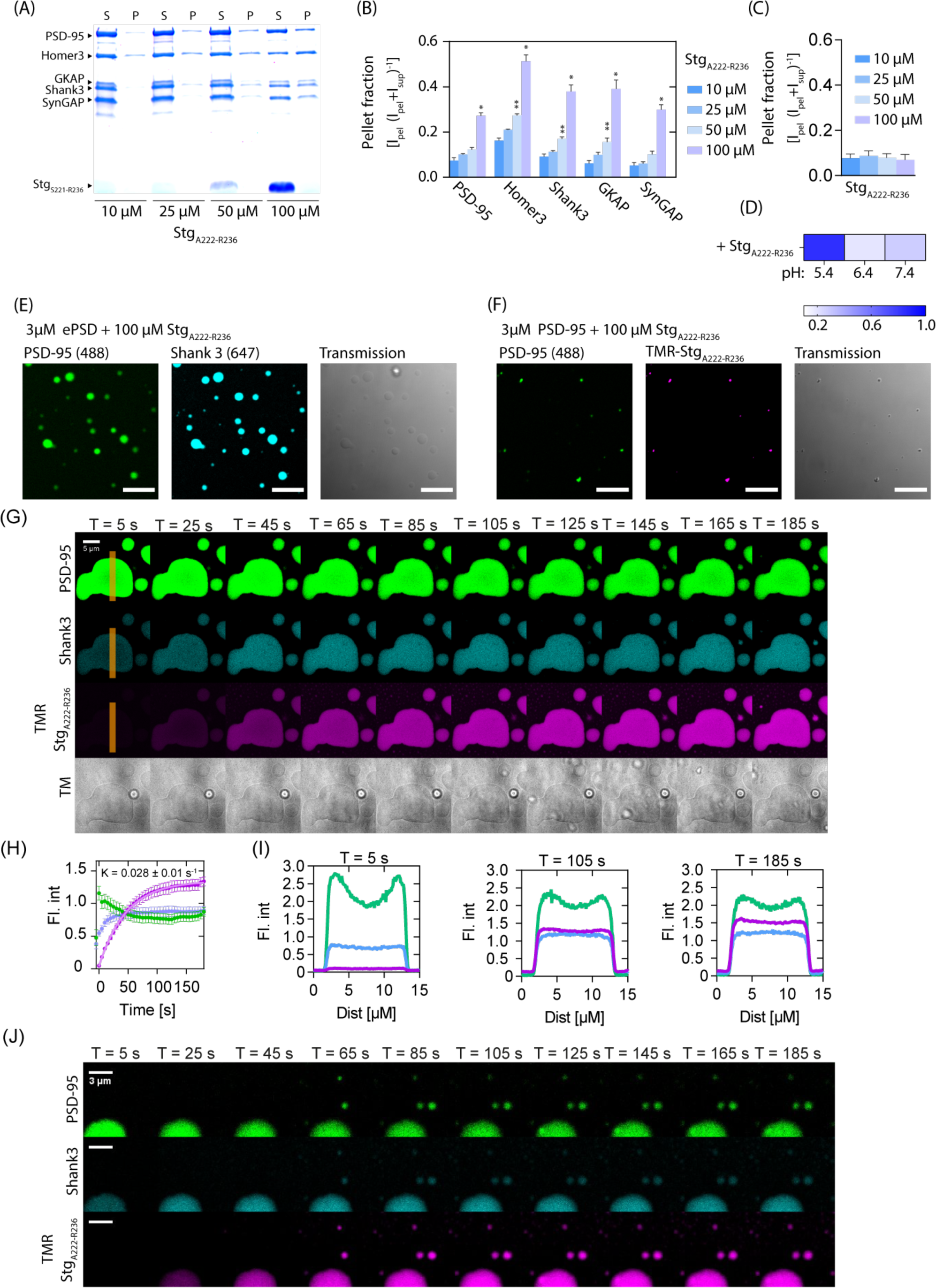
Stg_A222-R236_ is sufficient to induce LLPS. (A-B) SDS-PAGE sedimentation assay with ePSD (3 µM) incubated with increasing amounts of Stg_A222-R236_, show increasing LLPS. (A) SDS-PAGE sedimentation assay with 5xePSD (3 µM) incubated with increasing amounts of Stg_A222-R236_. (B) Quantification of the gels shown in (A) shows increased tendency for LLPS formation with increasing concentrations of Stg_A222-R236_. (C) Quantification of SDS-PAGE sedimentation assay with PSD-95 (10 µM) incubated with increasing amounts of Stg_A222-R236_. (D) pH titration of PSD-95 in presence of Stg_A222-R236_ shows increased LLPS formation at lower pH. (E-F) Images validating that Stg_A222-R236_ can induce LLPS for the ePSD (D) but not for PSD-95 (E) alone. (G) Time series of H-S-G-S condensate first incubated with PSD-95 and subsequent addition of TMR labelled Stg_A222-R236_, shows uptake into existing droplets. (H) Quantification of mean droplet fluorescence intensity shows rapid uptake of TMR Stg_A222-R236_. Error bars are shown as SEM of 17 droplets, fitting was done using GraphPad Prism using a single exponential association fit. (I) Line scans of droplet intensity for PSD-95 (green), Shank (blue), and Stg_A222-R236_ (purple). (J) Time series of H-S-G-S condensate first incubated with PSD-95 and subsequent addition of TMR labeled Stg_A222-R236_, shows formation of new droplets.

In addition to the integration into existing droplets, Stg_A222-R236_ increased the LLPS formation as evidenced by appearance of new, smaller, droplets over the time of the experiment (Figure 6J). Besides Stg_A222-R236_, we also generated a 32-mer peptide covering the entire predicted interaction region of the A214-E245 region (Stg_A214-E245_) (Figure S10). The SDS-PAGE sedimentation assay showed similar results for Stg_A214-E245_ as for Stg_A222-R236_, however, fluorescence confocal microscopy showed that when Stg_A214-E245_ was mixed with the ePSD the mixture progressed towards a pH dependent (not shown) meso-crystal like precipitate (Figure S10), similar to literature examples when LLPS systems progress from the highly dynamic LLPS droplets to less dynamic aggregates and fibrils (Ray et al., 2020). FRAP showed that the Stg_A214-E245_ induced precipitate remained dynamic (Figure S10).

The above sedimentation assays and imaging demonstrated that a 15-mer peptide comprising residues A222-R236 derived from the Stg C-terminal, while inadequate for formation of LLPS with PSD-95 alone, is sufficient to promote LLPS condensate formation for the ePSD and reorganize the dynamic network.

### Known inhibitors of the PSD-95 PDZ domains can influence LLPS

Several inhibitors have been developed to target the PDZ domains of PSD-95, including two promising drug candidates that feature Arg-rich cell penetrating peptides (CPPs) (Bach et al., 2012; Christensen et al., 2019; Aarts et al., 2002). To potentially bridge the mechanistic understanding between LLPS and drug discovery, we wanted to investigate whether the two clinically relevant peptide inhibitors of PSD-95, NA-1 (nerinetide) and AVLX-144, affected LLPS formation for either PSD-95 or the ePSD. NA-1 is a monomeric 20-mer peptide containing the 9 C-terminal amino acids from the GluN2B NMDAR subunit, fused to an N-terminal 11-mer peptide from the trans-activator of transcription protein (TAT), which facilitates cell penetration of NA-1. The clinical investigations of NA-1 have provided the compound in a dosage of 2.6 mg/kg, which corresponds approximately to 1 µM the compound distributed in the entire body volume. AVLX-144 is likewise, derived from the GluN2B C-terminal, but is a bivalent inhibitor, also comprising a TAT peptide. Both NA-1 and AVLX-144 targets the first two PDZ domains of PSD-95 through canonical PDZ interactions (Bach et al., 2012; Aarts et al., 2002) and both adopt a random coil structure in solution (Figure S11A).

We found that NA-1 (36 µM) induced LLPS when incubated with Alexa633-PSD-95 (30 nM labelled/3 µM unlabeled) at pH below 7.4 (Figure 7A), while AVLX-144 only induced LLPS at pH below 6.4 (Figure 7B). This was validated using SDS-PAGE sedimentation (Figure 7C and Figure S11B-C). To show that the pH change did not compromise the affinity of NA-1 towards PSD-95, we did competitive FP. At pH 5.4, we saw an inverted competition curve (Figure 7D), suggesting formation of larger molecular assemblies as a function of NA-1 concentration (K_i,app_, pH 5.4 = 156 µM, SEM interval: [155-158] µM). This suggests that NA-1 can induce LLPS at a more acidic pH of pH 5.4, but importantly also at very low PSD-95 concentrations ([CPSD-95] =150 nM). Interestingly we observed only minor changes in K_i_ as a function of pH (Figure 7D), in the pH range 6.4-9.4 (K_i,app, pH 6.4_ = 7.2 µM, SEM interval: [6.0-8.4] µM; K_i,app, pH 7.4_ = 4.5 µM, SEM interval: [3.4-5.7] µM; K_i,app, pH 8.4_ = 2.6 µM, SEM interval: [1.5-3.8] µM; K_i,app, pH 9.4_ = 4.1 µM, SEM interval: [2.9-5.2] µM), this was also the case for saturation binding of 5FAM-labelled dimeric inhibitor to PSD-95 (Figure S11D).

**Figure 7.**
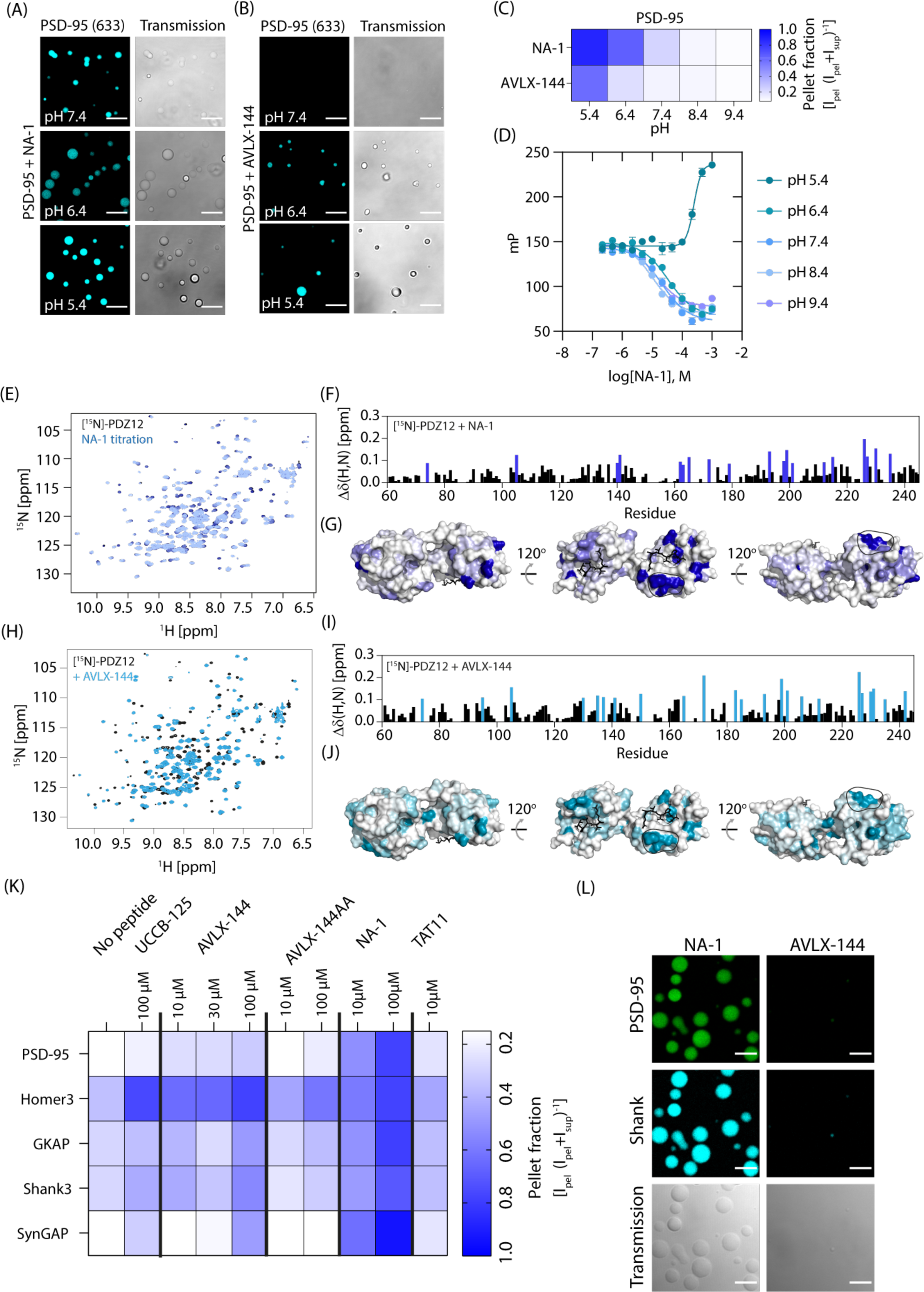
Pharmacological inhibitors of PSD-95 affect LLPS formation for PSD-95 and ePSD. (A-B) Confocal microscopy shows that NA-1 (A) and AVLX-144 (B) and induces LLPS once complex with PSD-95 at indicated pH values. (C) Heat map representation of SDS-PAGE sedimentation quantification of NA-1 and AVLX-144 pH dependent LLPS induction. (D) Competitive FP at different pH suggests drastic changes in the complex size at low pH, as indicated by the upward sigmoidal curve, compared to the downward sigmoidal curves for higher pH values. (E) [^1^H]-[^15^N]-HSQC spectra overlay of 100 μM PDZ1-2 titrated with NA-1 peptide concentrations ranging from 4 μM to 500 μM. (F) Chemical shift perturbation on PDZ1-2 (PDB: 3GSL) upon addition of NA-1 show wide spread perturbations in the PDZ1-2 tandem. Blue bars indicates perturbations larger than the mean Δδ(H,N) + 1 Std. Dev. (G) Surface representation of PDZ1-2 with NA-1 inducted perturbations larger than the mean Δδ(H,N) + 1 Std. Dev. Highlighted in blue, while remaining Δδ(H,N) were colored with a gradient from blue to white according to their Δδ(H,N). The black ligands represent RTTPV, which was docked into the PDZ binding pocket of both PDZ1 and PDZ2 using alignment to PDB ID 3JXT (Sainlos et al., 2011). (H) [^1^H]-[^15^N]-HSQC spectra overlay of free 100 μM PDZ1-2 (black) and with 512 μM AVLX-144 peptide and PDZ1-2 (teal). (I) Chemical shift perturbation on PDZ1-2 upon addition of AVLX-144 show widespread perturbations in the PDZ1-2 tandem. Teal bars indicates perturbations larger than the mean Δδ(H,N) + 1 Std. Dev. (J) Surface representation of PDZ1-2 (PDB: 3GSL) with AVLX-144 induced perturbations larger than the mean Δδ(H,N) + 1 Std. Dev. Highlighted as teal, while remaining Δδ(H,N) were colored with a gradient from teal to white according to their Δδ(H,N). The black ligands represent RTTPV, which was docked into the PDZ binding pocket of both PDZ1 and PDZ2 using alignment to PDB ID 3JXT. (K) Heatmap representation of SDS-PAGE sedimentation quantification of indicated peptides (see Figure S10 gels and quantification). (L) Confocal microscopy shows that NA-1, but not AVLX-144, induces LLPS once complexed with the ePSD.

To obtain molecular insight into the mechanism underlying the differential propensity of NA-1 and AVLX-144 to cause LPPS, we recorded ^1^H-^15^N-HSQC NMR experiments of PDZ1-2 with and without these peptides (Figure 7E-J). Both peptides caused similar changes of the chemical shifts throughout the primary sequence of PDZ1-2 suggesting comparable binding for both NA-1 and AVLX-144. Both NA-1 and AVLX-144 perturbed residues in PDZ2 to a similar extent, while the perturbations in PDZ1 were stronger for AVLX-144 (Figure 7F-G) than for NA-1 (Figure 7I-J), in concordance with dual occupancy of both PDZ domains in the PDZ1-2 tandem for AVLX-144 (Bach et al., 2012; Chi et al., 2010; Aarts et al., 2002). We suspected that the positively charged TAT peptide in NA-1 was able to utilize the negatively charged patch on PDZ1, as have been suggested to be the secondary binding site for the S/R rich region of Stg (Zeng et al., 2019), but we did not see any perturbations indicating this, for neither NA-1 nor AVLX-144. We also tested the 11-mer TAT peptide alone and Stg_A222-R236_, which did not result in any chemical shift perturbations in the recorded ^1^H-^15^N-HSQCs of PSD-95 (512 µM Stg_A222-R236_/128 µM ^15^N-PDZ1-2). The reason for the lack of pertubations might be that the isolated PDZ1 interactions, as presented earlier, occurs on a timescale which is outside of our experimental window.

### NA-1 can induce LLPS condensate formation in the ePSD complex

Expanding these findings into the ePSD system, we found that NA-1 could induce LLPS at 10 µM (* for PSD-95, p<0.0205; ns for Homer3, p=0.0947; ns for Shank3, p=0.0872; * for GKAP p=0.0252; * for SynGAP, p=0.0286; two-way ANOVA Dunnett post-test) and 100 µM (*** for PSD-95, p<0.001; ** for Homer3, p<0.01; * for Shank3, p=0.0280; ** for GKAP p<0.01; ** for SynGAP, p<0.01; two-way ANOVA Dunnett post-test) in the PSD-95 containing ePSD (Figure 7K and Figure S12). On the other hand, AVLX-144 induced LLPS to a minor extent (Figure 7E and Figure S12) at 10 µM (* for PSD, p=0.0190; ns for Homer3, p=0.185; ns for Shank3, p=0.910; ns for GKAP, p=0.993; ns for SynGAP, p=0.148; two-way ANOVA Dunnett post-test) and 100 µM (* for PSD, p=0.0357; ns for Homer3, p=0.117; ns for Shank3, p=0.814; ns for GKAP, p=0.993; ns for SynGAP, p=0.247; two-way ANOVA Dunnett post-test). Mutation of the PBM of AVLX-144 (TDV to ADA) to generate AVLX-144-AA, a non-binding version, largely compromised the ability to drive LLPS of the ePSD (Figure S12). The TAT peptide alone (10 µM) induced LLPS (** for PSD-95, p<0.01; ns for Homer3, p=0.05; * for Shank3, p=0.0189; * for GKAP p=0.0252; ns for SynGAP, p=0.0949; two-way ANOVA Dunnett post-test) of the ePSD similar to AVLX-144 (Figure 7K and Figure S12). Removal of the TAT peptide from AVLX-144, generating AVLX-125 (UCCB-125), did not alter the ePSD LLPS formation (Figure 7K). AVLX-125 (100 µM) pellet formation was only significant for Homer3 and SynGAP (ns for PSD-95, p=0.45; * for Homer3, p<0.05; ns for Shank3, p=0.87; ns for GKAP p>0.99; ** for SynGAP, p=0.0026; two-way ANOVA Dunnett post-test) of the ePSD similar to AVLX-144 (Figure 7K and Figure S12G). Taken together this suggests a novel mechanism for peptide inhibitors of PPIs, highlighting inhibitors as LLPS drivers in PPI networks, indicating that seemingly inert parts of the peptide for the specific protein interaction, can facilitate more flexible protein interaction networks, thus suggesting a novel way of targeting protein-protein interactions.

## DISCUSSION

The recent discovery of phase separation of key PSD components including PSD-95 reveals a highly complex phenomenon and provides a new paradigm for the understanding of synaptic biology. Since it was shown that proteins massively concentrate in the condensate droplets, LLPS have emerged as a possible explanation of how the postsynapse can be immensely enriched in proteins, and how these are sorted solely on the basis of their protein interaction networks (Alberti et al., 2019; Feng et al., 2019a; Yoshizawa et al., 2020; Zeng et al., 2018; Zeng et al., 2019; Zeng et al., 2016a).

We found that PSD-95 adopts a compact structure in solution, suggesting intramolecular interactions, which is in good accordance with previous studies. As shown recently PDZ1-2 can undergo spontaneous oligomerization (Rodzli et al., 2020), a fact that we also observed when increasing the PDZ1-2 concentration to the mM range. Under normal circumstances protein concentrations in the mM range is considered non-physiological, but due to recent developments in cell biology, mainly focused on LLPS (Feng et al., 2019a; Feng et al., 2019b; Yoshizawa et al., 2020), it seems increasingly common that protein complexes which undergo LLPS are of high µM and even mM concentration in the LLPS droplets (Feng et al., 2019a; Feng et al., 2019b; Yoshizawa et al., 2020), suggesting that even low affinity and low specificity interactions becomes of high importance in these supracomplexes.

The ability of PSD-95 to take part in LLPS has recently been described in great detail in pioneering work by Zhang and co-workers (Feng et al., 2019b; Tao et al., 2019; Zeng et al., 2019; Zeng et al., 2016a). We observed that Shank3, Homer3, GKAP and SynGAP together undergo LLPS at low concentration and that PSD-95 acts as a negative regulator for LLPS when mixed with the remaining components. This occurs in absence of a PSD-95 PDZ1-2 specific ligand. PSD-95 has earlier been shown to facilitate LLPS in the ePSD system, but in earlier cases this negative regulatory effect has not been observed. Earlier work conducted on the ePSD system (Zeng et al., 2018; Zeng et al., 2019) shows that PSD-95 does not actively participate in LLPS in the absence of a multivalent ligand, a tetrameric construct of the GluN2B C-terminal, also using the GCN4p1 backbone (Zeng et al., 2018), however based on our experiments using the multivalent Stg constructs, and previous publications (Zeng et al., 2018; Zeng et al., 2019; Zeng et al., 2016a), it is evident that PSD-95 can undergo separate LLPS, which subsequently can incorporate into the Homer/Shank/GKAP LLPS system, through low affinity GKAP linkage (K_D_ = 176 µM (Zeng et al., 2018)).

It is known that low and moderate affinity interactions can push systems towards LLPS, as is the case here for both the isolated Stg_A222-R236_ peptide and the PSD-95 inhibitor, NA-1. This has also been shown earlier for intrinsically disorder proteins, which in some cases can act as promiscuous LLPS drivers (Protter et al., 2018). The ability to drive LLPS has also shown earlier that Arg rich peptides can induce LLPS in large protein sets (Boeynaems et al., 2017).

NA-1 and AVLX-144 are both promising clinical drug candidates in the treatment of acute ischemic stroke (Bach et al., 2019; Hill et al., 2020). In our experiments we were able to show that NA-1 induced LLPS upon interaction with isolated PSD-95 and also in the more complex setting of the ePSD, in which case the concentration of NA-1 that induces LLPS is close to the clinically relevant concentration of NA-1. This suggests that the TAT peptide used in both NA-1 and ALVX-144 can in some cases bind to a different part of PSD-95, not being the PDZ1-2, which is not possible for AVLX-144, potentially due to the high affinity of AVLX-144, which offers less interaction flexibility than NA-1. The ability of known pharmacologically relevant peptides and the cell penetrating peptide TAT to induce LLPS uncovered here potentially points to a novel molecular mechanism for some of the neuroprotective effects of these compounds, and in therapeutically relevant concentrations. The difference in binding for NA-1 and AVLX-144 suggests that AVLX-144 is more tightly bound to both PDZ1-2 while NA-1, as expected, favors PDZ2 binding, but is also able to bind PDZ1. The absence of perturbations to the acidic residues in PDZ1, suggests that the residues responsible for TAT induced LLPS is positioned elsewhere in PSD-95. It will be interesting to see if the ability to induce LLPS in the very simple ePSD system translates into functional effects in the treatment of acute ischemic stroke or similar diseases which relies on the dynamic functions of the PSD.

## ACKNOWLEDGEMENTS

We would like to thank Prof. Mingjie Zhang and Prof. Daniel Choquet for providing plasmids. We would also like to thank Simon Erlendsson for fruitful discussions and Nabeela Khadim for technical assistance. We also acknowledge the Core Facility for Integrated Microscopy, Faculty of Health and Medical Sciences, University of Copenhagen and the cOpenNMR facility (Novo Nordisk Foundation, NNF18OC0032996) at Department of Biology, University of Copenhagen. We gratefully acknowledge SAXS beamtime at the P12 beamline at the PETRAIII at DESY, Germany along with help from the beamline scientist. We also kindly acknowledge the team at FIDABio (Copenhagen, Denmark) for their great help and assistance. D.E.O. thanks the Carlsberg Foundation (grant nr. CF14-0284) and the Novo Nordisk Foundation (grant nr. 12953) for support.

## AUTHOR CONTRIBUTIONS

N.R.C, K.S and K.L.M conceived the project. N.R.C performed protein expression and purification, conducted and analyzed data from circular dichroism, size exclusion chromatography, fluorescence polarization, flow induced dispersion analysis, SDS-PAGE sedimentation assays, fluorescence confocal microscopy and celluSPOT experiments. N.R.C and L.A performed and analyzed the small angle X-ray scattering experiments. C.P.P and K.T.E performed and analyzed the NMR experiments. N.R.C., J.N.P and D.E.O. performed and analyzed the size exclusion chromatography multi angle light scattering experiments. A.T.S and N.R.C designed multimeric peptides. V.S designed, synthesized and printed the celluSPOT arrays. M.V.M did LC-MS and UPLC. N.R.C wrote the manuscript with contributions from all authors.

## CONTACT FOR REAGENT AND RESOURCE SHARING

Information and request for sharing of reagents, plasmids etc. should be directed to Lead contacts, Kristian Strømgaard or Kenneth Lindegaard Madsen

## METHODS DETAILS

### Plasmid preparation

Plasmids encoding FL-PSD-95 (32M3C-PSD-95 FL), ΔN-PSD-95 (32M3C-PSD-95 61-724), Homer3 (M3C-Homer 3 EVH1-CC WT), Shank3 (M3C-Shank3 NPDZ-HBS-CBS-SAM M1718E), GKAP (32M3C-GKAP 3GBR-CT) and SynGAP (MG3C-SynGAP CC-PBM WT) was a kind gift from Prof. Mingjie. Zhang (The Division of Life Science, Hong Kong University of Science and Technology). In brief, all constructs were previously cloned into a pET32a containing an N-terminal thioredoxin (TRX) tag or a streptococcal protein G (GB1) tag followed by a 6xHis affinity tag and a Prescission C3 protease site, followed by the protein of interest as described in (Zeng et al., 2018; Zeng et al., 2019; Zeng et al., 2016a).

Stargazin AA variant was prepared using the QuickChange site directed mutagenesis kit (Agilent Technologies, USA), using stargazin WT (A kind gift from D. Choquet, IINS, Bordeaux, France) as the template and the following primers, 3’-CCGCCGGACC**G**CCCCCG**CC**TAAGGATCCGAAGGGC-5’ and 5’-GCCCTTCGGATCCTTA**GG**CGGGGG**C**GGTCCGGCGG-3’.

DNA encoding rat Arc 195-364 (Uniprot: Q63053) was ordered from ThermoFischer with an 3’-BamHI and a EcoRI site followed by a FactorXa protease site and the Arc 195-364 coding sequence, followed by a 5’HindIII, NotI and XhoI cleavage site. The plasmid was inset into a pGEX4T1 vector using the BamHI and XhoI sites, resulting in a construct encoding an N-terminal GST followed by a thrombin and a factorXa protease site followed by rat Arc 195-364.

### Recombinant protein expression and purification

Plasmids encoding FL-PSD-95 (32M3C-PSD-95 FL 1-724), ΔN-PSD-95 (32M3C-PSD-95 61-724), Homer3 (M3C-Homer 3 EVH1-CC WT), Shank3 (M3C-Shank3 NPDZ-HBS-CBS-SAM M1718E), GKAP (32M3C-GKAP 3GBR-CT), SynGAP (MG3C-SynGAP CC-PBM WT) and GST-Arc 195-364 were grown in BL21-DE3-pLysS *E. coli* in LB medium supplemented with 100 µg/mL ampicillin (HelloBio, #HB4322) and 25 µg/mL chloramphenicol (Sigma, C0378). The growth was induced at OD_600_ 0.6-0.8 with 0.5-1 mM IPTG (Sigma 10724815001) and cultures were grown for 16 hrs at 18°C shaking at 170 rpm. Cells were harvested at 7000 g and frozen at -80 °C until purification. Pelleted cells were suspended in lysis buffer containing 50 mM Tris (Sigma 93362) (pH 8.0), 300 mM NaCl (S9888), 1 mM TCEP (Sigma C4706), half a tablet of cOmplete™ Protease Inhibitor Cocktail (Sigma 11697498001) and 2.5 µg/mL Deoxyribonuclease I (Sigma D5025) and sonicated (Branson Sonifier 250, 3 mm round tip, 40% output, 70/30 pulse) on ice until solution became homogeneous. Lysate was centrifuged at 36.000 g for 30 min at 20 °C and supernatant was collected and purified using affinity chromatography. 6xHis proteins were purified using a HisTrap HP 5 mL column (GE Life science 17524701) using an imidazole gradient from 10-500 mM imidazole (Sigma 56749). GST-Arc was purified by addition of 700 µL/L culture Glutathione Sepharose 4B beads (GE Life Science, 17075605) in lysis buffer. The GST beads were washed using centrifugation (4000 g for 5 min at 25 °C) followed by removal of supernatant and addition of 50 mM Tris (pH 7.4), 300 mM NaCl, 10 mM EDTA, 1 mM TCEP, this step was repeated twice, followed by transfer of beads to a single use gravity column (BioRad 7326008), followed by three on column washes. The protein was eluted using 10 mM reduced GSH (Sigma G4251) in 50 mM Tris (pH 7.4), 300 mM NaCl, 10 mM EDTA, 1mM TCEP. In case of both 6xHis and GST tagged protein, the affinity chromatography was followed by buffer change and purification using size exclusion chromatography (HiLoad 16/600 Superdex 200 pg, GE Life science 28989335) in 50 mM Tris (pH 8.0), 300 mM NaCl, 10 mM EDTA, 1 mM TCEP. Mass and purity was validated using LC-MS and UPLC to be >93% for all purified proteins where concentrated to a suitable concentration, aliquoted and flash frozen in liquid nitrogen. Before fluorescence labeling and use in assays protein was exchanged into PBS-TECP (PBS-TCEP) using NAP-5 columns (GE Life science #17085301), pre-equilibrated in PBS-TCEP.

### Protein labeling

Before labelling 1 mg of solid dye (NHS-AlexaFlour647, ThermoFischer A20006; NHS-AlexaFlour568, ThermoFischer A20103; C5 Maleimide-AlexaFluor633, ThermoFischer A20342; AlexaFluor488 C5 maleimide) was diluted into 10µL DMSO (Sigma #D2650) and aliquoted (0.02 mg/tube) and DMSO was evaporated using vacuum evaporation, aliquots were stored at -20 °C until usage. For protein labeling, dyes, were dissolved in DMSO and purified protein in PBS-TCEP was incubate with respective dye for 1-2 h. For NHS reactions, the reaction was quenched by addition of 100 mM Tris (pH 7.4). Excess dye and Tris was removed using two consecutive NAP5 columns equilibrated with PBS-TCEP. Protein and dye concentration was measured by NanoDrop 3000 (ThermoFischer). For confocal imaging fluorescent protein was diluted to a final ratio of 1/10 with unlabeled protein.

### Peptide array synthesis

μSPOT peptide arrays (CelluSpots, Intavis AG, Cologne, Germany) were synthesized using a ResPepSL synthesizer (Intavis AG) on acid labile, amino functionalized, cellulose membrane discs (Intavis AG) containing 9-fluorenylmethyloxycarbonyl-β-alanine (Fmoc-β-Ala) linkers (minimum loading 1.0 μmol/cm). Synthesis was initiated by Fmoc deprotection using 20% piperidine in *N*-methylpyrrolidone (NMP) (1 × 2 and 1 × 4 μL, 3 and 5 min, respectively) followed by washing with dimethylformamide (DMF, 7 × 100 μL per disc) and ethanol (EtOH, 3 × 300 μL per disc). Peptide chain elongation was achieved using 1.2 μL of coupling solution consisting of preactivated amino acids (0.5 M) with 2-(1-benzotriazole-1-yl)-1,1,3,3-tetramethyluronium hexafluorophosphate (0.5 M) and *N*,*N*-diisopropylethylamine (DIPEA) in NMP (2:1:1, amino acid:HBTU:DIPEA). The couplings were carried out 7 times (20 min for the first coupling and 30 min for the rest), and subsequently, the membrane was capped twice with capping mixture (5% acidic anhydride in NMP), followed by washes with DMF (7 × 100 μL per disc). After chain elongation, final Fmoc deprotection was performed with 20% piperidine in NMP (3 × 4 μL, 5 min each), followed by 6 washes with DMF, subsequent N-terminal acetylation with capping mixture (3 × 4 μL, 5 min each) and final washes with DMF (7 × 100 μL per disc) and EtOH (7 × 200 μL per disc). Dried cellulose membrane discs were transferred to 96 deep-well blocks and were treated with the side-chain deprotection solution consisting of 80% trifluoracetic acid (TFA), 12% DCM, 5% H_2_O, and 3% triisopropylsilane (TIPS) (150 μL per well) for 1.5 h at room temperature. Afterwards, the deprotection solution was removed, and the discs were solubilized overnight at room temperature using a solvation mixture containing 88.5% TFA, 4% trifluoromethansufonic acid (TFMSA), 5% H_2_O, and 2.5% TIPS (250 μL per well). The resulting peptide-cellulose conjugates were precipitated with ice-cold diethyl ether (1 mL per well) and spun down at 1000 rpm for 90 min, followed by an additional wash of the formed pellet with ice-cold diethyl ether. The resulting pellets were re-dissolved in dimethyl sulfoxide (DMSO, 500 μL per well) to give final stocks, which were transferred to a 384-well plate and printed (in duplicates) on white coated CelluSpots blank slides (76 × 26 mm, Intavis AG) using a SlideSpotter robot (Intavis AG).

### Peptide synthesis

Purified (>95% purity) TAT11 (YGRKKRRQRRR), mono-Stg (biotin-ahx-RMKQLEPKVEELLPKNYHLENEVARLKKLVGGGGSRRTTPV), dim-Stg (biotin-ahx-RMKQLEDKVEELLSKNYHLENEVARLKKLVGGGGSRRTTPV), tri-Stg (biotin-ahx-RIKQIEDKIEEILSKIYHIENEIARIKKLIGGGGSRRTTPV) were ordered and from TAGCopenhagen (Denmark). Purified (>95% purity) AVLX-144 was ordered from WuXi peptides (China). UCCB-125 and AVLX-144-AA were synthesized in house using previously reported synthesis (Bach et al., 2009; Bach et al., 2012)

The synthesis of the Stg_A222-R236_ peptide (GAITRIPSYRYRYQRR), using Fmoc-based solid phase peptide synthesis, was carried on a Prelude X, induction heating assisted, peptide synthesizer (Gyros Protein Technologies, Tucson, AZ, USA) with 10 mL glass reaction vessel using preloaded Wang-resins (100–200 mesh). All reagents were prepared as solutions in DMF: Fmoc-protected amino acids (0.2 M), O-(1H-6-chlorobenzotriazole-1-yl)-1,1,3,3-tetramethyluronium hexafluorophosphate (HCTU, 0.5 M) and DIPEA, 1.0 M. Coupling steps were carried out using the following protocol: deprotection (20% piperidine in DMF, 2 × 2 min, room temperature, 300 rpm shaking), coupling (2 × 5 min, 75 °C, 300 rpm shaking, for Arg and His couplings 2 × 5 min, 50 °C, 300 rpm shaking). Amino acids were double coupled using amino acid/HCTU/DIPEA (ratio 1:1.25:2.5) in 5-fold excess over the resin loading to achieve peptide sequence elongation.

N-terminal labeling of peptide A222-R236 with 5 (and 6)-carboxytetramethylrhodamine (TAMRA, Anaspec Inc.) was performed on resin, by coupling TAMRA for 16 h at room temperature using a mixture of 1.5:1.5:3 [TAMRA:benzotriazol-1-yloxy)tripyrrolidinophosphonium hexafluorophosphate (PyBOP): DIPEA] in NMP (Witte et al., 2013). The coupling was finalized with extensive washes of resin with DMF and DCM. The synthesized peptides were cleaved from the resin using a mixture of 90:2.5:2.5:2.5:2.5 (TFA:H_2_O:TIPS:1,2-ethanedithiol (EDT):thioanisole) for 2 h at room temperature. After cleavage the peptide was precipitated with an ice-cold diethyl ether and centrifuged at 3500 rpm for 10 min at 4 °C. The resulting peptide precipitate was re-dissolved in 50:50:0.1 (H_2_O:CH_3_CN:TFA) and lyophilized. Purification of the crude peptide was performed with a preparative reverse phase high performance liquid chromatography (RP-HPLC) system (Waters) equipped with a reverse phase C18 column (Zorbax, 300 SB-C18, 21.2 × 250 mm) and using a linear gradient with a binary buffer system of H_2_O:CH_3_CN:TFA (A: 95:5:0.1; B: 5:95:0.1) (flow rate 20 mL/min). The collected fractions were characterized by LC-MS. The purity (≥95%) of the fractions was determined at 214 nm on RP-UPLC. The final lyophilized products were used in further experiments.

### Size exclusion chromatography

Before analytical SEC, stocks in PBS-TECP were mixed to desired concentrations in PBS- TECP and incubated for 20 min at room temperature before being run on a Superdex 200 increase 10/300 gl column (GE Lifescience 28990944) monitoring the Absorbance at 220 nm, 260 nm and 280 nm. The resulting absorbance trace at 260 (peptides alone) 280 nm (in complex with PSD-95) was plotted and normalized to the maximal absorbance for each condition. For peptide and protein containing samples, the data was normalized to the maximal absorbance of the protein sample in absence of peptide. Data was plotted using GraphPad Prism 8.3.

### SEC Multi Angle Light Scattering (MALS)

SEC-MALS was done using an Agilent HPLC equipped with a Wyatt MALS setup, where 50 µL of 50 µM PSD-95 incubated with 150 µM of indicated peptide was loaded onto a Superdex200 Increase 10/300 column equilibrated in 50 mM Tris (pH 7.5), 200 mM NaCl, 1 mM TCEP and both absorbance, refractive index and light scattering data was collected. Resulting data was analyzed and molecular weight was calculated using the ASTRA® software package, data was plotted using GraphPad Prism 8.3.

### Small-angle X-ray scattering (SAXS)

Concentration series of FL-PSD95 ranging from 30-210 µM was prepared in buffer containing 50 mM Tris (pH 8.2), 300 mM NaCl, 5 mM EDTA, 1 mM TCEP. Samples were measured at the P12 SAXS Beamline, PetraIII, DESY, Hamburg Germany (Blanchet et al., 2015). Preliminary data reduction including radial averaging and conversion of the data into absolute scaled scattering intensity, I(q), as a function of the scattering vector q, where q= 4π sin(θ)/λ (θ = half scattering angle, λ = wavelength) were done using the standard procedures at the beamline (Blanchet et al., 2015). Scattering data was merged; buffer subtracted and binned using WillItRebin with a binning factor of 1.02. *p*(*r*) functions were fitted using Bayesian Indirect Fourier transformations (GenApp.Rocks). Ensemble optimization method was run using known structural domains of PSD-95 (PDZ1-2, PDB: 3GSL; PDZ3, PDB: 5JXB; SH3-GK, PDB: 1KJW) as structural domains and linkers were modeled as fully-flexible linkers. EOM (Bernado et al., 2007; Tria et al., 2015) (Ranch) was used to generate 10.000 structural models with 15 harmonics, and 10% symmetric structures. EOM (Gajoe) was used to fit the pool (10.000 models). Gajoe was run using 1000 generations in the genetic algorithm with 100 ensembles, a non-fixed ensemble size, with a maximum of 20 curves pr. ensemble and 100 repetitions. The fit was evaluated on the basis of their χ^2^ value and their *D_max_* and *R_g_* distributions.

### Flow Induced dispersion analysis (FIDA)

FIDA was carried out using intrinsic fluorescence, using the standard protocol recommended by the manufacturer, in short, PSD-95 (12 µM) in absence or presence of 12 µM or 36 µM peptide was loaded to the FIDA1 instrument, the protein containing solution was used as injectant and the protein with peptide was used as analyte solution. The diffusion of the complex could then be observed using intrinsic fluorescence, and the hydrodynamic radius was calculated using the FIDA software 2.0 using a single Gaussian distribution fit, at 75% and curve smoothing. Resulting hydrodynamic radius was plotted using GraphPad Prism 8.3.

### Fluorescence Polarization

Fluorescence Polarization (FP) saturation binding (also described in (Bach et al., 2012; Madsen et al., 2005)) was carried out in a buffer containing 50 mM Tris (pH 8.0), 300 mM NaCl, 10 mM EDTA, 1 mM TCEP, using an increasing amount of protein incubated with a fixed concentration of fluorescently labeled peptides as indicated. Competition FP was done at a fixed concentration of PSD-95 and a bivalent fluorescent tracer, AB-143 (Bach et al., 2012), against an increasing concentration of unlabeled peptide. After mixing the 96-well plate (a black half-area Corning Black non-binding) was incubated 20 min on ice after which the fluorescence polarization was measured directly on a Omega POLARstar plate reader using excitation filter at 488 nm and long pass emission filter at 535 nm. The data was plotted using GraphPad Prism 8.3 and fitted to the either a single exponential binding curve or a sigmoidal single site binding model for saturation experiments or One site competition for competition experiments. K_i_’s were automatically calculated using the Cheng-Prusoff equation. All binding isotherms were repeated at least 3 technical replicates or as indicated in figure legend.

### Circular Dichroism (CD)

Before CD measurements samples were diluted into 50 mM NaPi buffer (pH 8.0) to a suitable concentration, Stg peptides 8 µM was used and for NA-1 and AVLX-144 10 µM was used. CD measurements were done using a Jasco J1500 at 25 °C with a quartz cell with a 1 mm path length quartz cuvette. Each spectrum was recorded from 260-190 nm at a 0.1 nm step resolution and a scan speed of 50-100 nm/min, each presented spectrum is the average of three scans. The resulting mDEG signal was converted into molar ellipticity, θ (deg ⋅ cm^2^ ⋅ dmol) using the equation θ = (mDEG·10^6^)/(C·N·L), where mDEG is the measured signal, C is the protein concentration in µM, N is the number of residues in the protein, L is the cuvette path length in mm. The resulting CD spectra were plotted using GraphPad Prism 8.3.

### SDS-PAGE sedimentation assay

Proteins were all mixed in the desired concentration in PBS-TCEP and equilibrated for 10 min before centrifugation at 20 000 g for 15 min at 25 °C using a temperature-controlled table top centrifuge. Following centrifugation, the supernatant was collected and the pellet was re-suspended in an equal amount of PBS-TCEP, usually 50 µL. To ensure proper suspension of LLPS samples were vortexed before addition of SDS buffer boiling at 95°C for 5 min. Supernatant and pellet fractions were run on any kD™ Mini-PROTEAN^®^ TGX™ Precast Protein Gels (10 or 15 wells, BioRad 4569036 or 4569033). Gels were imaged using a Li-COR Odyssey gel scanner and band intensities were analyzed using ImageJ. Significance was evaluated using one-way ANOVA with Dunnett post-test, one-way ANOVA with Tukey post-test or a two-way ANOVA with Dunnett post-test.

### Confocal microscopy on LLPS droplets

Confocal microscopy was done using a Zeiss LSM780 using a 63x NA 1.4 plan apochromat oil objective using Argon 488 nm 25 mW, 543 nm HeNe 1.2 mW and 633 nm HeNe 5mW lasers using a detection wavelength of 490-538 nm for the 488 channel, 556-627 nm for the 543 channel, 636-758 for the 633 channel. Images were acquired using averaging of 4 line scans and 12-bit. The LLPS droplets were prepared in the desired concentration in PBS-TCEP at desired pH, mostly pH 7.4 unless stated otherwise, and added to an untreated lab tec (155411PK, Nunc^TM^, ThermoFischer) and imaged after being allowed to settle for 15 min at 25°C. For samples containing fluorescent protein or peptide the content of fluorescent protein or peptide was kept at 1-10% of indicated total protein or peptide concentration. Fluorescence after photo bleaching (FRAP) experiments was done by bleaching of the 488nm or 647 nm channel, normalizing the fluorescence intensity to ROI intensity before bleaching to 1 and immediately after bleaching to 0.

### NMR spectroscopy

All NMR spectra were recorded on 600 MHz or 750 MHz Bruker Avance III HD spectrometers equipped with QCI or TCI cryo-probes at 25 °C in 50 mM Tris pH 8, 200 mM NaCl, 1 mM TCEP, 10 % D2O and 250 µM DSS. Spectra were processed with NMRPipe (Delaglio et al., 1995) or qMDD (Kazimierczuk and Orekhov, 2011) if non-uniform sampling was used for the acquisition and analyzed using CCPNMR analysis (Skinner et al., 2016). Amide nitrogen and proton chemical shift assignments were kindly shared by Prof. Mingjie Zhang and validated using HNCA and HN(CO)CA experiments on a sample with 300 µM ^13^C^15^N PSD-95 PDZ1-2. Ligand titrations were followed by ^1^H-^15^N-HSQC experiments recorded on 100 µM ^15^N PSD-95 and ligand concentrations ranging from 500 µM to 4 µM. Combined chemical shift perturbations were calculated between the unbound and the bound states using: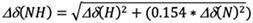

## DATA AVAILABILITY

All primary data and custom scripts are available upon request from the corresponding authors.

