## Supplementary material for "Bi-directional protein-protein interactions control liquid-liquid phase separation of PSD-95 and its interaction partners": SI

### SUPPLEMENTAL INFORMATION TITLES AND LEGENDS:

### SUPPLEMENTARY FIGURES AND TABLES:

#### Supplementary figures

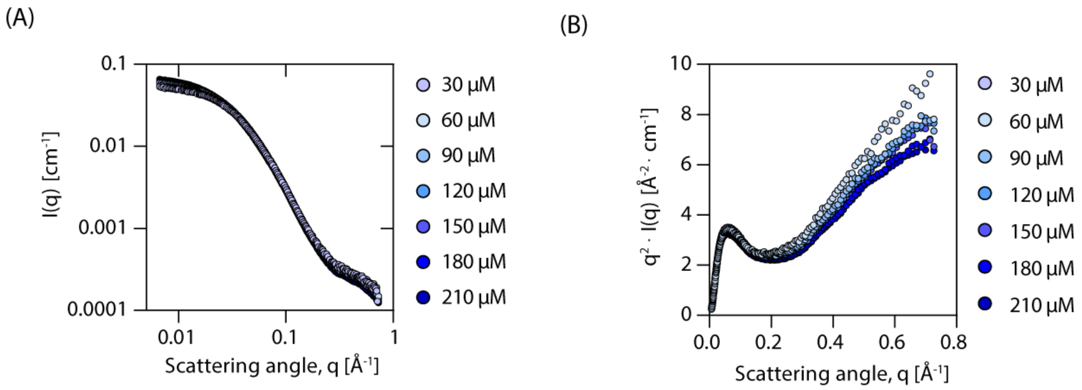
(A)
0.1
0.01
0.001
0.0001
I(q) [cm⁻¹]
0.01
0.1
1
Scattering angle, q [Å⁻¹]
30 µM
60 µM
90 µM
120 µM
150 µM
180 µM
210 µM
(B)
10
8
6
4
2
0
q² · I(q) [Å⁻² · cm⁻¹]
0.0
0.2
0.4
0.6
0.8
Scattering angle, q [Å⁻¹]
30 µM
60 µM
90 µM
120 µM
150 µM
180 µM
210 µM

**Figure S1** (A) SAXS scattering profiles of PSD-95 in the concentration range from 30-210 μM. (B) Kratky representation of (A) indicates that PSD-95 is a multi-domain protein containing flexible linkers. (B) Also show excluded volume repulsion, as seen from the drop in (q)/c at low q-values, at the highest concentrations of PSD-95, as is to be expected for high protein concentrations. For this reason we focused our analysis on the SAXS data obtained at 60 μM.

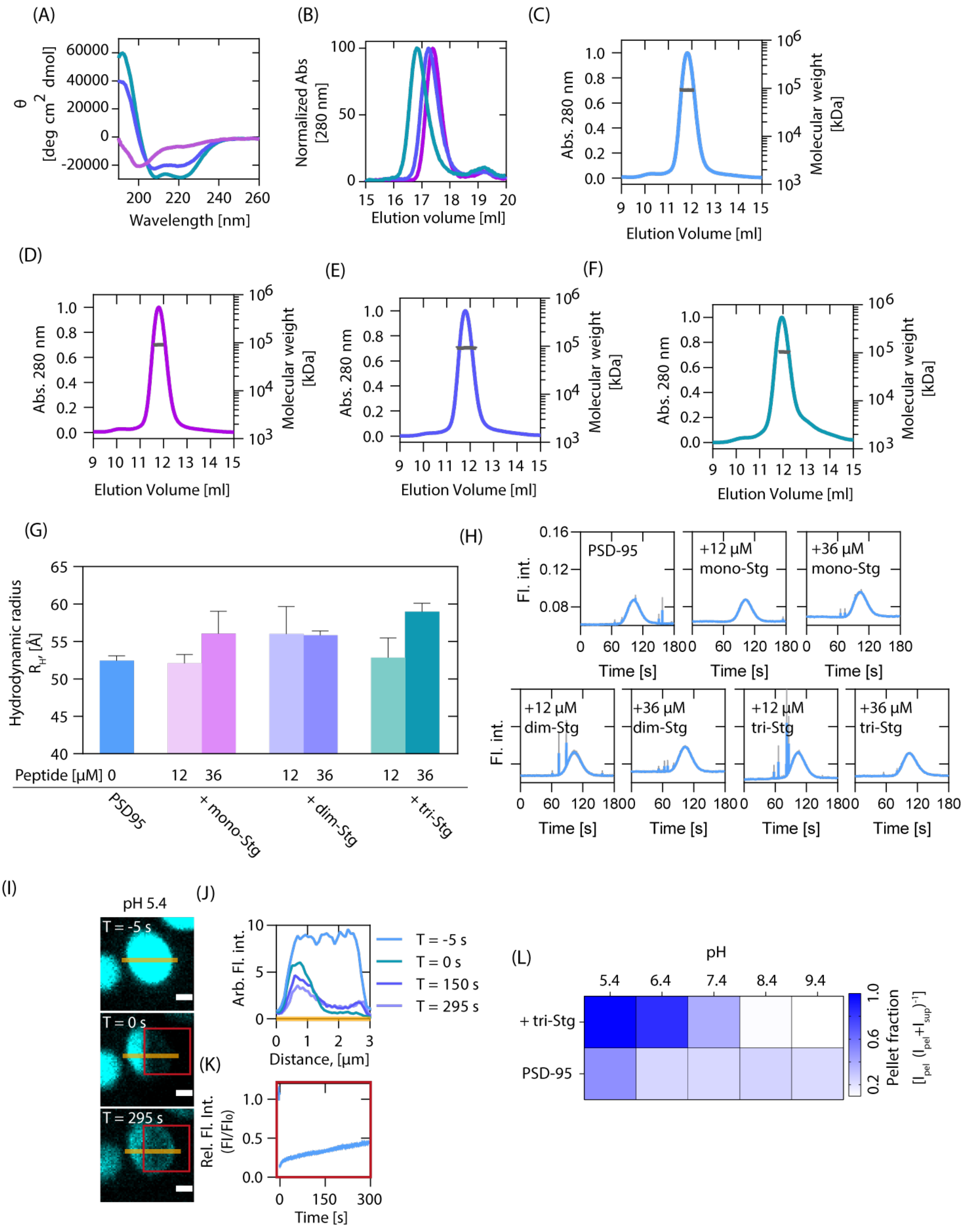
(A)
θ [deg cm2 dmol]
60000
40000
20000
0
-20000
200 220 240 260
Wavelength [nm]
(B)
Normalized Abs [280 nm]
100
50
0
15 16 17 18 19 20
Elution volume [ml]
(C)
Abs. 280 nm
Molecular weight [kDa]
Elution Volume [ml]
(D)
Abs. 280 nm
Molecular weight [kDa]
Elution Volume [ml]
(E)
Abs. 280 nm
Molecular weight [kDa]
Elution Volume [ml]
(F)
Abs. 280 nm
Molecular weight [kDa]
Elution Volume [ml]
(G)
Hydrodynamic radius $R_H$, [Å]
Peptide [μM] 0 12 36 12 36 12 36
PSD95
+ mono-Stg
+ dim-Stg
+ tri-Stg
(H)
Fl. int.
PSD-95
+12 μM mono-Stg
+36 μM mono-Stg
+12 μM dim-Stg
+36 μM dim-Stg
+12 μM tri-Stg
+36 μM tri-Stg
Time [s]
(I)
pH 5.4
T = -5 s
T = 0 s
T = 295 s
(J)
Arb. Fl. int.
T = -5 s
T = 0 s
T = 150 s
T = 295 s
Distance, [μm]
(K)
Rel. Fl. Int. (Fl/Fl0)
Time [s]
(L)
pH
5.4 6.4 7.4 8.4 9.4
+ tri-Stg
PSD-95
Pellet fraction $[I_{pel} (I_{pel}+I_{sup})^{-1}]$

**Figure S2.** (A) Circular dichroism of mono-Stg (purple), dim-Stg (blur) and tri-Stg (green) shows helical structure for dim-Stg and tri-Stg, while mono-Stg adopts a random coil like structure. (B) Size exclusion chromatography of mono-Stg (purple), dim-Stg (blue) and tri-Stg (green) indicates an increase in hydrodynamic radius for tri-Stg over mono-Stg and dim-Stg.(C-F) SEC-MALS elution profiles of 50 μM TRX-PSD-95 in absence (C) or presence of 150 μM mono-Stg (D), dim-Stg (E) or tri-Stg (F) and fitted molecular weights. (G) FIDA obtained hydrodynamic radius of 12 μM TRX-PSD-95 (blue) in complex with mono-Stg (purple), dim-Stg (blue) or tri-Stg (green) indicates a slight increase in size for the complex, as also indicated from SEC-MALS data (C-F), errorbars are shown as SEM of N=3. (H) Average FIDA taylorgrams of (J), show spikes in signal possibly due to LLPS droplets in some mixtures. Error is shown in grey interval as SEM of N=3. (I) Heat-map representation of SDS-PAGE sedimentation assay with full length PSD-95 in absence or presence of tri-Stg indicates strong pH dependency of multivalent LLPS formation. (J) Representative FRAP image time series of LLPS droplet PSD-95 at pH 5.4. (K)  Line intensity profile of PSD-95 at indicated timepoints, which show a time dependent reduction in PSD-95 and minor recovery of PSD-95 signal. (L) Quantification of PSD-95 droplets FRAP recovery at pH 5.4 suggest some dynamics in the PSD-95 droplets.

1163

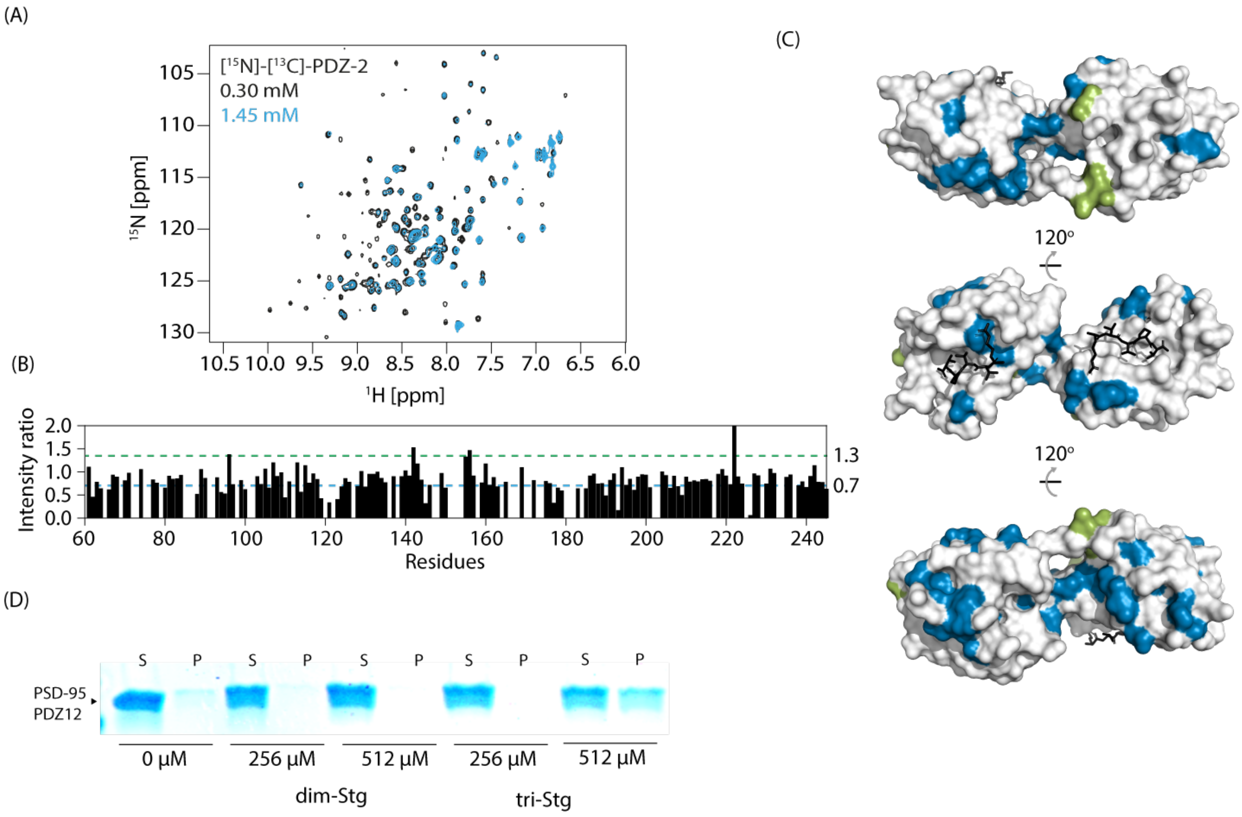
(A)
[15N]-[13C]-PDZ-2
0.30 mM
1.45 mM
15N [ppm]
1H [ppm]
(B)
Intensity ratio
Residues
1.3
0.7
(C)
120°
120°
(D)
S P S P S P S P S P
PSD-95 PDZ12
0 µM
256 µM
512 µM
256 µM
512 µM
dim-Stg
tri-Stg

**Figure S3** (A) $^{1}$H-$^{15}$N-HSQC spectra overlay of $^{15}$N-labelled PSD-95 PDZ12 at 1.45 mM (teal) and 300 μM (black). (B) Peak intensity ratios of $^{1}$H-$^{15}$N-HSQC spectra in (B), where peaks were normalized according to concentrations and the ratio was taken as $I_{peak,1.45\ mM}$ / $I_{peak,0.3\ mM}$. (C) Residues with intensity ratio change over 30%, was mapped onto the structure of PDZ1-2 (PDB: 3GSL), with intensity decrease over 30% (<0.7 in (B)) are presented in teal and intensity increases over 30% (>1.3 in (B)) are presented in green and black ligands represents RTTPV which was docked into the PDZ binding pocket of both PDZ1 and PDZ2 using alignment to PDB ID 3JXT. (D) SDS-PAGE sedimentation assay of PDZ1-2 in absence or presence of indicated peptide.

1173

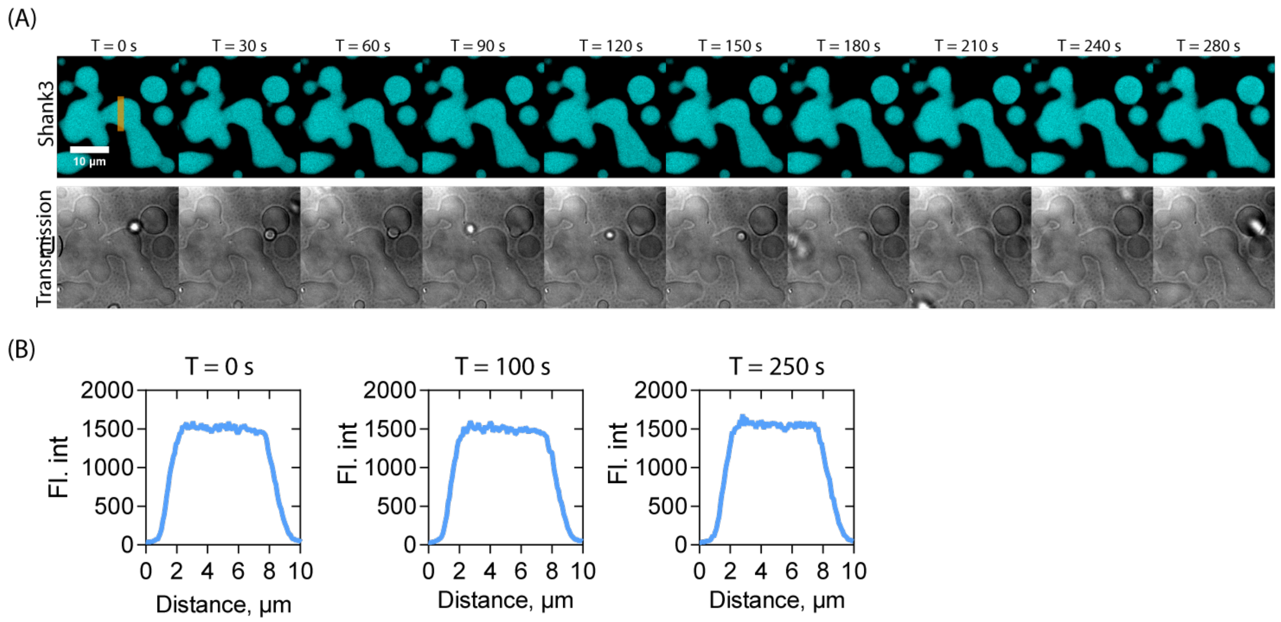


1174

1175

**Figure S4** (A) Time series of H-S-G-S condensate after addition of PBS. (B) Line profile of Shank3 at indicated time points shows no difference in Shank3 intensity upon addition of PBS. Orange line in (A) indicates line segment.

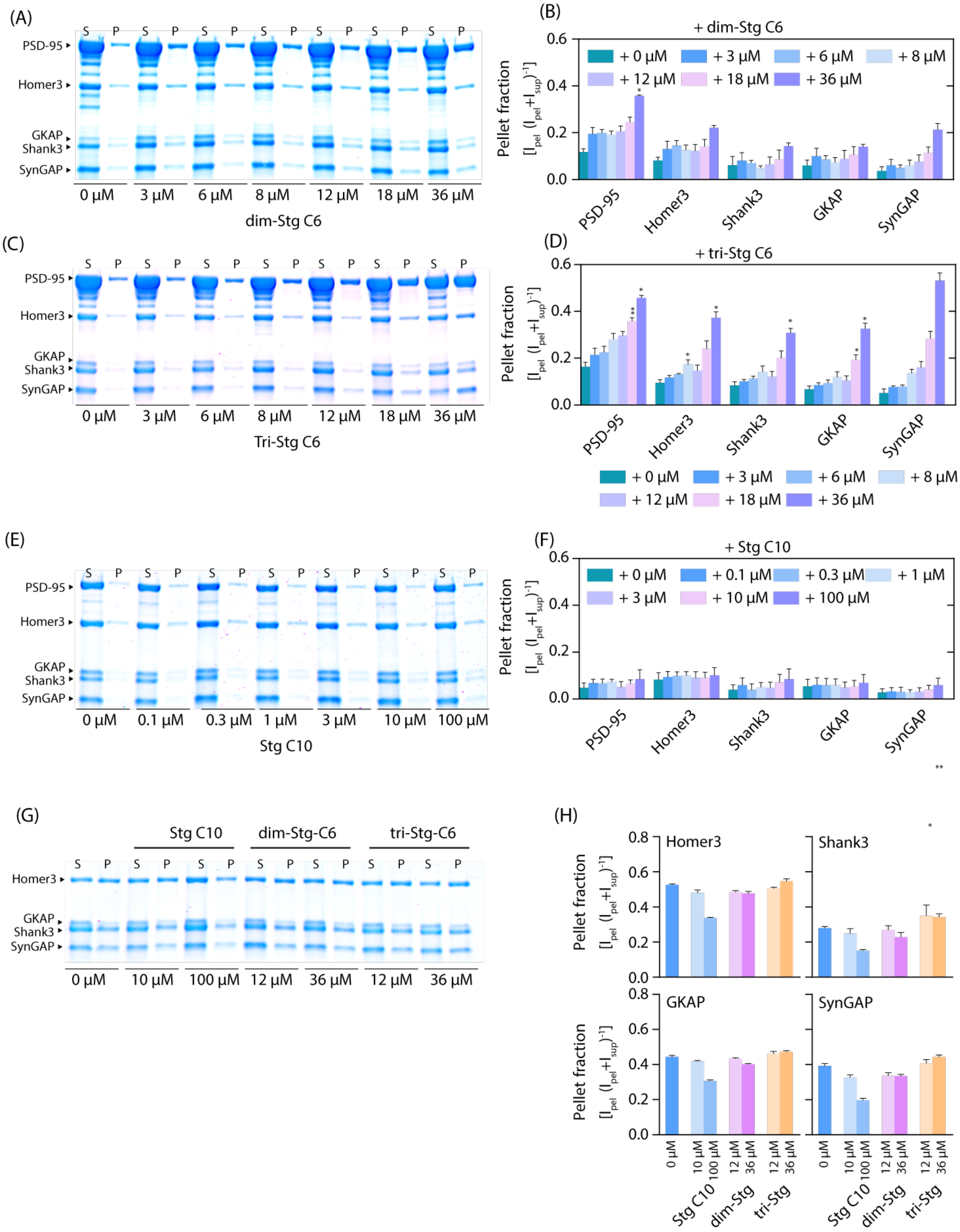
(A)
S P S P S P S P S P S P S P
PSD-95
Homer3
GKAP
Shank3
SynGAP
0 μM 3 μM 6 μM 8 μM 12 μM 18 μM 36 μM
dim-Stg C6
(B)
+ dim-Stg C6
+ 0 μM + 3 μM + 6 μM + 8 μM
+ 12 μM + 18 μM + 36 μM
Pellet fraction
PSD-95 Homer3 Shank3 GKAP SynGAP
(C)
0 μM 3 μM 6 μM 8 μM 12 μM 18 μM 36 μM
Tri-Stg C6
(D)
+ tri-Stg C6
(E)
0 μM 0.1 μM 0.3 μM 1 μM 3 μM 10 μM 100 μM
Stg C10
(F)
+ Stg C10
+ 0 μM + 0.1 μM + 0.3 μM + 1 μM
+ 3 μM + 10 μM + 100 μM
(G)
Stg C10 dim-Stg-C6 tri-Stg-C6
0 μM 10 μM 100 μM 12 μM 36 μM 12 μM 36 μM
(H)
Homer3
Shank3
GKAP
SynGAP
Stg C10 dim-Stg tri-Stg

**Figure S5** (A-D) SDS-PAGE sedimentation and quantification of 5xePSD (3 μM H-S-G-S, 10 μM PSD-95) incubated with increasing amounts of dim-Stg (A-B) or tri-Stg (C-D). (E-F) SDS-PAGE sedimentation and quantification of 5xePSD (3 μM H-S-G-S, 3 μM PSD-95) incubated with increasing amounts of StgC10. Heatmap representation of A-D is shown in Figure 4H-I. (G) SDS-PAGE sedimentation assay with 3 μM H-S-G-S condensate incubated with indicated amount of Stg derived peptides. (H) Quantification of (G) shows no effects of peptide addition on condensate formation. Error bars are shown as SEM of n=3.

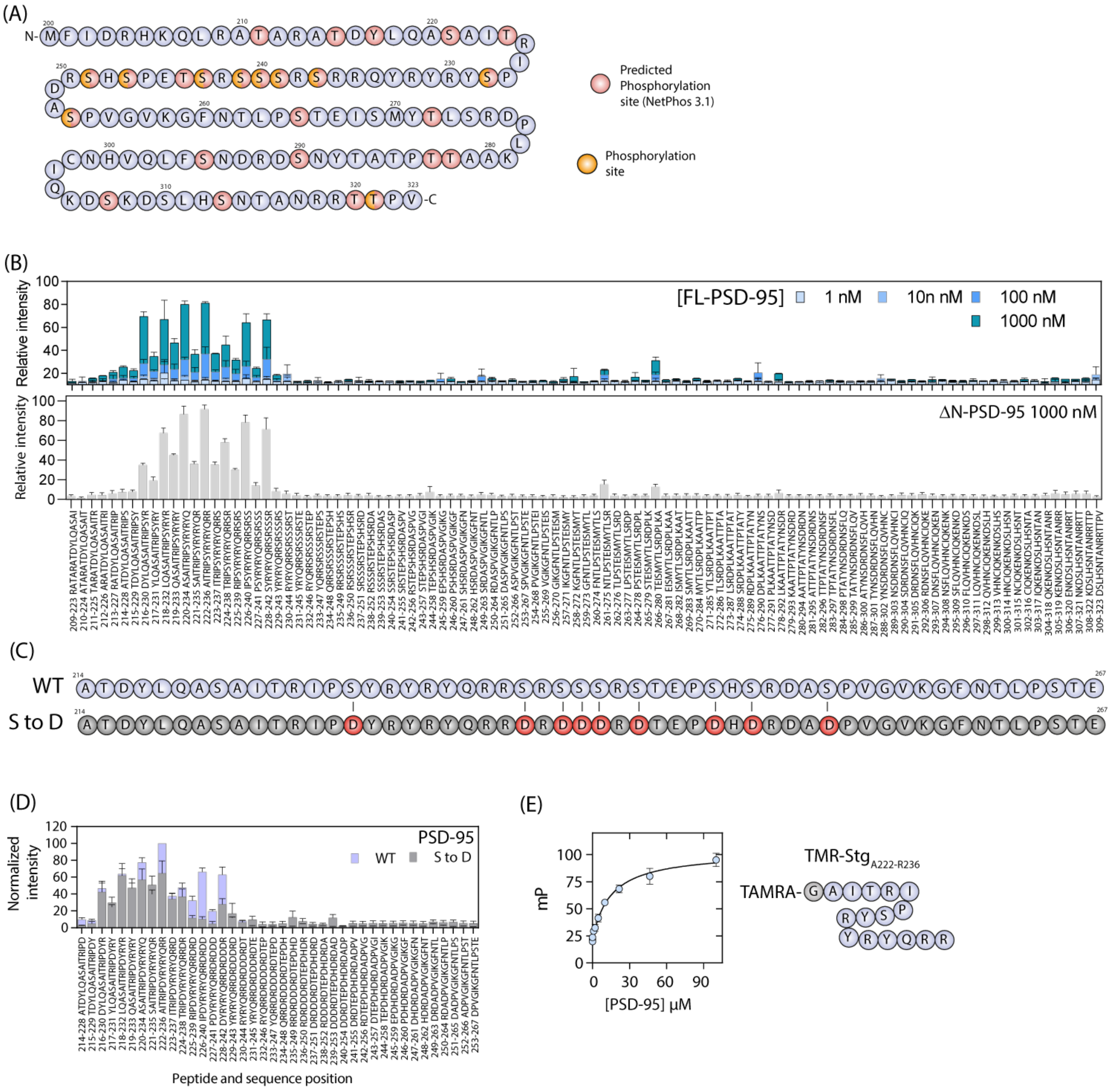
(A)
Predicted Phosphorylation site (NetPhos 3.1)
Phosphorylation site
(B)
Relative intensity
[FL-PSD-95]
1 nM
10n nM
100 nM
1000 nM
ΔN-PSD-95 1000 nM
(C)
WT
S to D
(D)
Normalized intensity
PSD-95
WT
S to D
Peptide and sequence position
(E)
mP
[PSD-95] μM
TMR-Stg A222-R236
TAMRA-

**Figure S6** (A) Primary sequence of Stg cytoplasmic C-terminal (200-323) with known and predicted phosphorylation sites indicated. (B) Quantification of arrays of Stg C-terminal peptides (16-mers) when incubated with indicated protein. Primary sequence of peptides is indicated below each bar. Error bars are shown as SD of duplicate measurements. FL-PSD-95 and ΔN-PSD-95 was labelled with Alexa633. (C) Primary sequence with indicated Ser to Asp (S-to-D) mutations (red) mimicking the previously reported Ser phosphorylation. (D) Comparison between PSD-95 binding to WT (blue) and S-to-D (grey) peptide array. (E) FP saturation binding of TAMRA labelled $Stg_{A222\text{-}R236}$ to FL-PSD-95. Error bars are shown as SEM of n=3.

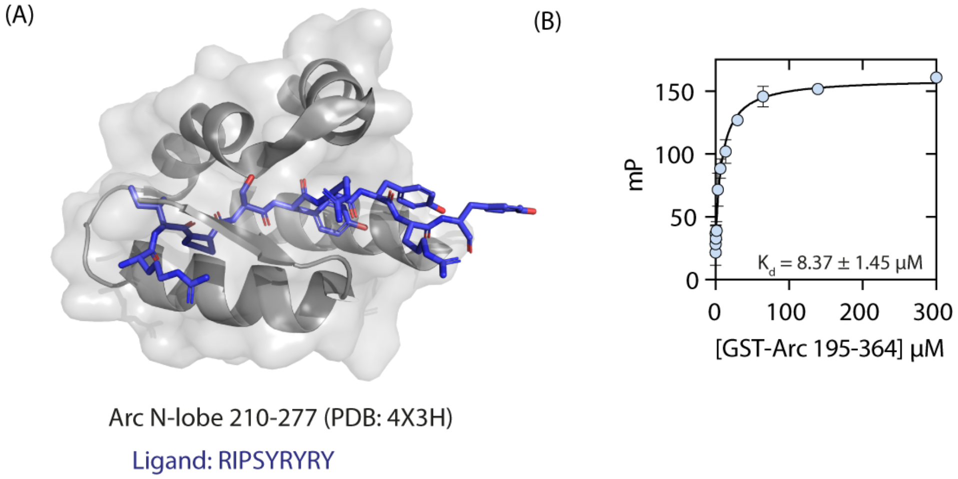
(A)
Arc N-lobe 210-277 (PDB: 4X3H)
Ligand: RIPSYRYRY
(B)
mP
150
100
50
0
0
100
200
300
K$_d$ = 8.37 ± 1.45 μM
[GST-Arc 195-364] μM

**Figure S7** (A) X-ray crystal structure (PDB: 4X3H) of Arc N-lobe (grey) binding to Stg peptide RIPSYRYRY (blue), (Zhang et al., 2015). (B) Fluorescence polarization binding of GST fused rat Arc N-/C-lob (Residues 195-364, Uniprot: Q63053). Fitting was done using GraphPad Prism 8.3, using a single binding site model. Error bars are shown as SEM of n=3.

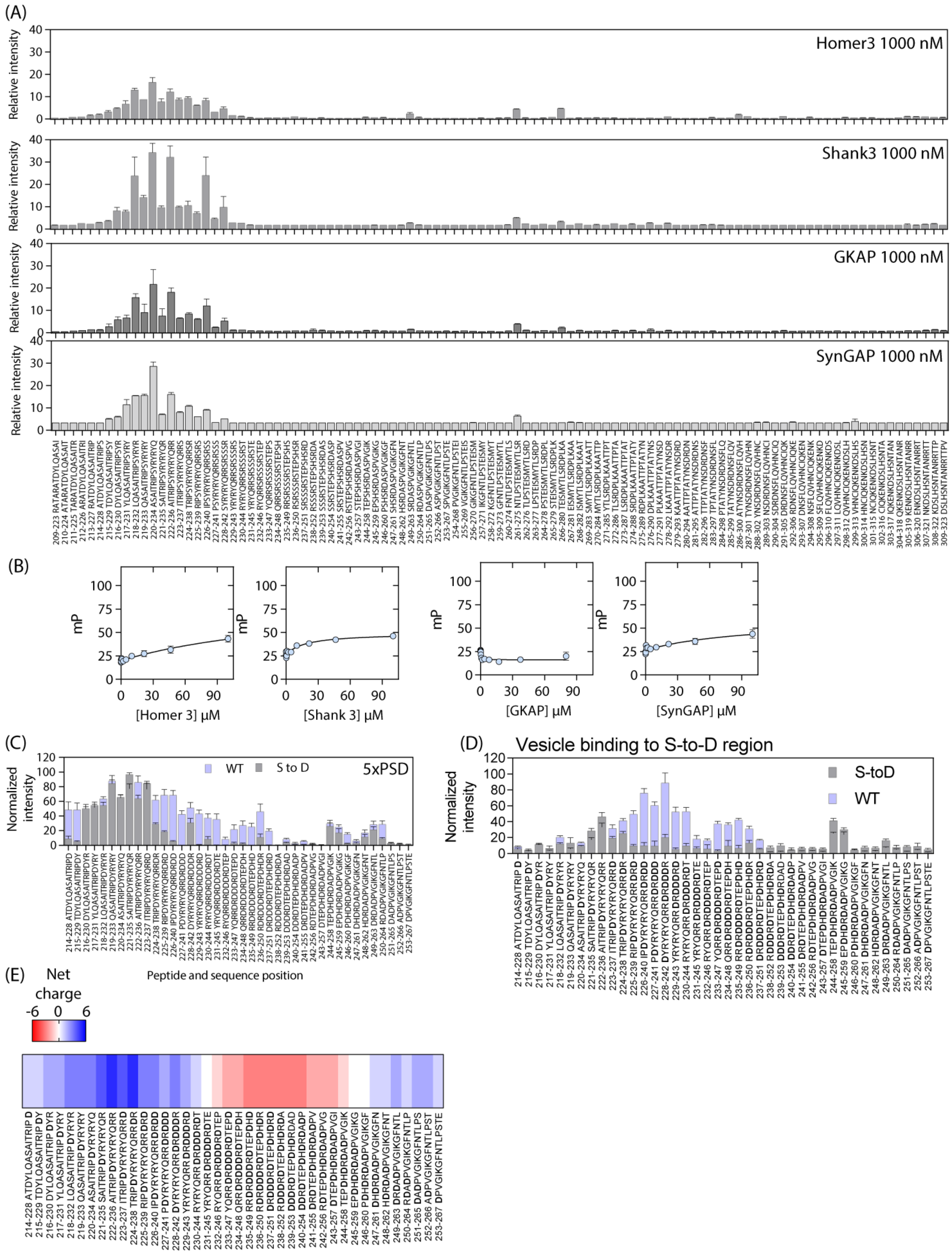
(A)
Homer3 1000 nM
Shank3 1000 nM
GKAP 1000 nM
SynGAP 1000 nM
Relative intensity
(B)
mP
[Homer 3] μM
[Shank 3] μM
[GKAP] μM
[SynGAP] μM
(C)
WT
S to D
5xPSD
Normalized intensity
Peptide and sequence position
(D)
Vesicle binding to S-to-D region
S-toD
WT
Normalized intensity
(E)
Net charge
-6
0
6

**Figure S8** (A) Quantification of arrays of Stg C-terminal peptides (16-mers) when incubated with indicated protein. Primary sequence of peptides is indicated below each bar. Error bars are shown as SD of duplicate measurements. Homer3, Shank3, GKAP and SynGAP were labelled with NHS-Alexa647. (B) Fluorescence polarization binding of TMR-$Stg_{A222\text{-}R236}$ to Homer3, Shank3, GKAP or SynGAP. Fitting was done using GraphPad Prism 8.3, using a single binding site model. Error bars are shown as SEM of n=3. (C) Comparison between ePSD binding to WT (blue) and S-to-D (grey) peptide array. (D) Binding to the S-to-D region for vesicles. Signal was normalized to maximal signal of WT peptides. Error bars are shown as SEM of n=6. WT data in is a regional crop from data shown in Figure 5C. (E) Net Charge distribution of S-to-D peptides.

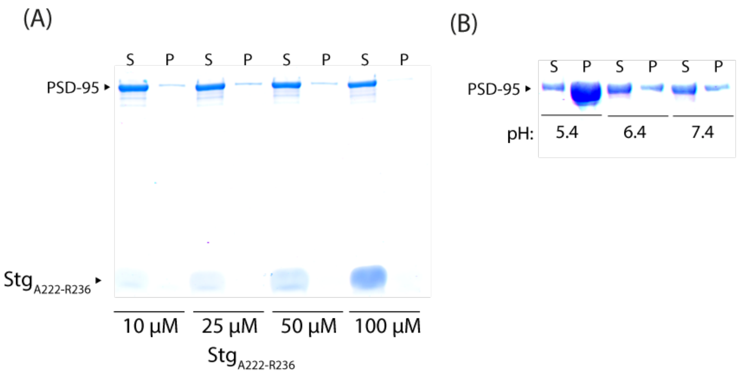


**Figure S9** (A) Representative SDS-PAGE sedimentation assay of PSD-95 incubated with StgA222-R236. For quantification see Figure 6C. (B) Representative SDS-PAGE sedimentation assay of PSD-95 incubated with 50 µM StgA222-R236 at different pH values.

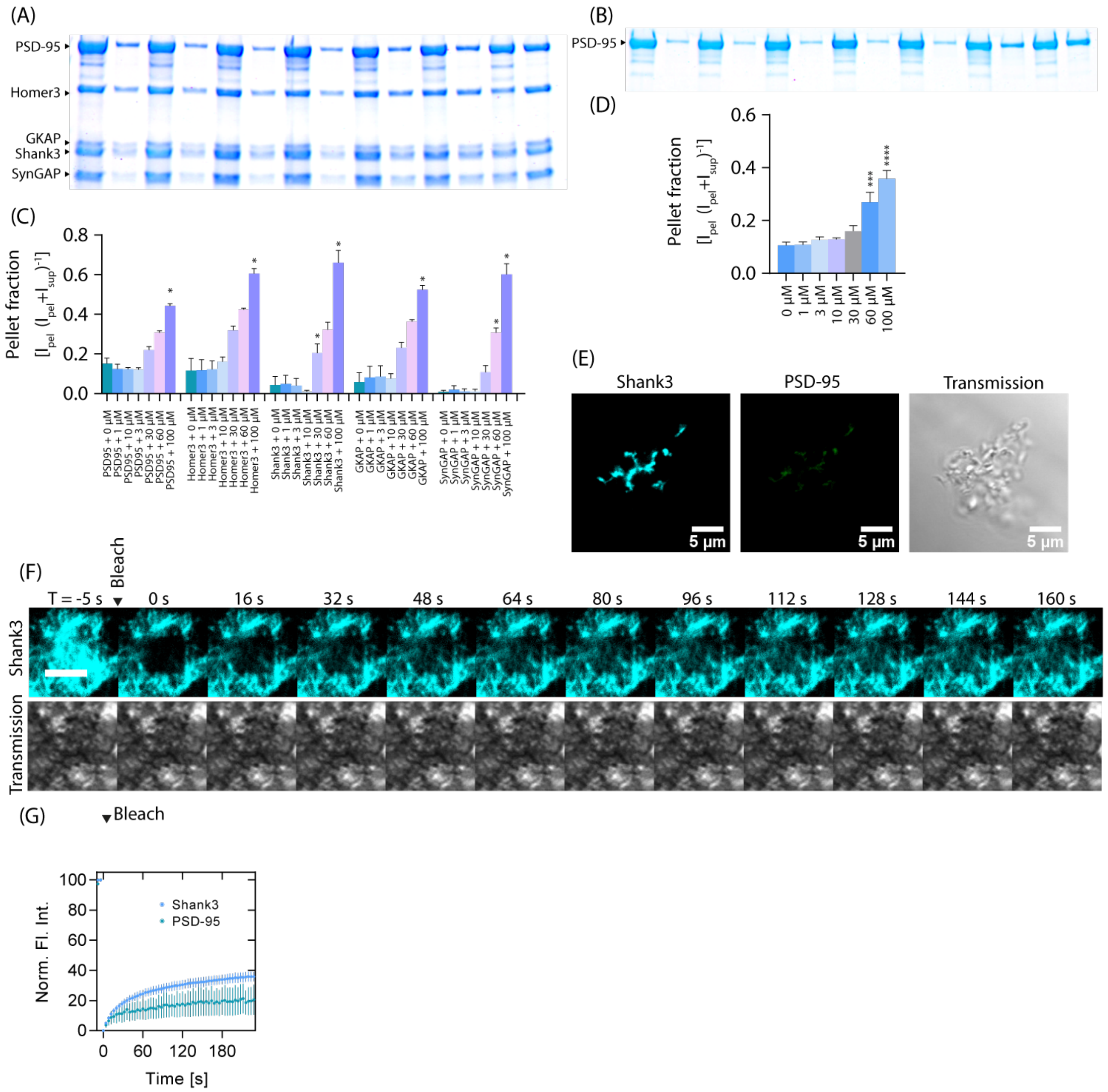


**Figure S10** (A-B) SDS-PAGE sedimentation assay of (A) ePSD or (B) PSD-95 with increasing amounts of $Stg_{A214\text{-}E245}$. (C-D) quantification of (A-B) shows a high degree of protein in the pellet fraction at concentrations above 30 µM $Stg_{A214\text{-}E245}$. Error bars shown as SEM of n=3. (E) Images of precipitate formed upon incubation of ePSD (3 µM) with 10 µM $Stg_{A214\text{-}E245}$. (F) FRAP image time series of 3 µM ePSD condensate incubated with 10 µM $Stg_{A214\text{-}E245}$ (G) Quantification of FRAP experiments suggests partial recovery of Shank3 intensity and minor recovery of PSD-95 intensity. Error bars are shown as SEM on n=3.

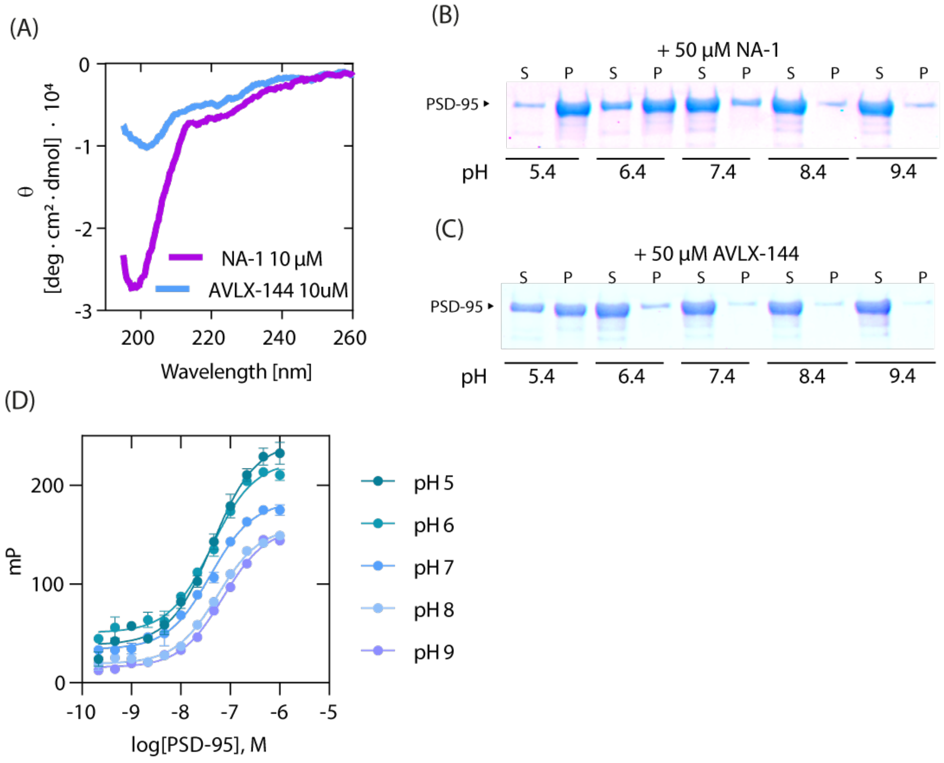

(A)
0
-1
-2
-3
θ
[deg · cm² · dmol] · 10⁴
NA-1 10 µM
AVLX-144 10uM
200
220
240
260
Wavelength [nm]
(B)
+ 50 µM NA-1
S P S P S P S P S P
PSD-95
pH
5.4
6.4
7.4
8.4
9.4
(C)
+ 50 µM AVLX-144
S P S P S P S P S P
PSD-95
pH
5.4
6.4
7.4
8.4
9.4
(D)
mP
200
100
0
-10
-9
-8
-7
-6
-5
log[PSD-95], M
pH 5
pH 6
pH 7
pH 8
pH 9

**Figure S11** (A) CD spectra of 10 μM NA-1 and AVLX-144 show that both are unstructured random coils. (B) Representative SDS-PAGE gel of 3 μM PSD-95 incubated with 50 μM NA-1. (C) Representative SDS-PAGE gel of 3 μM PSD-95 incubated with 50 μM AVLX-144. (D) FP saturation binding of 5 nM AB-143 (Bach et al., 2012) towards PSD-95 at different pH values, show no major change in KD, and changes in mP values are here accredited to change in fluorescent signal due to pH changes. Error bars indicate SEM of n=3.

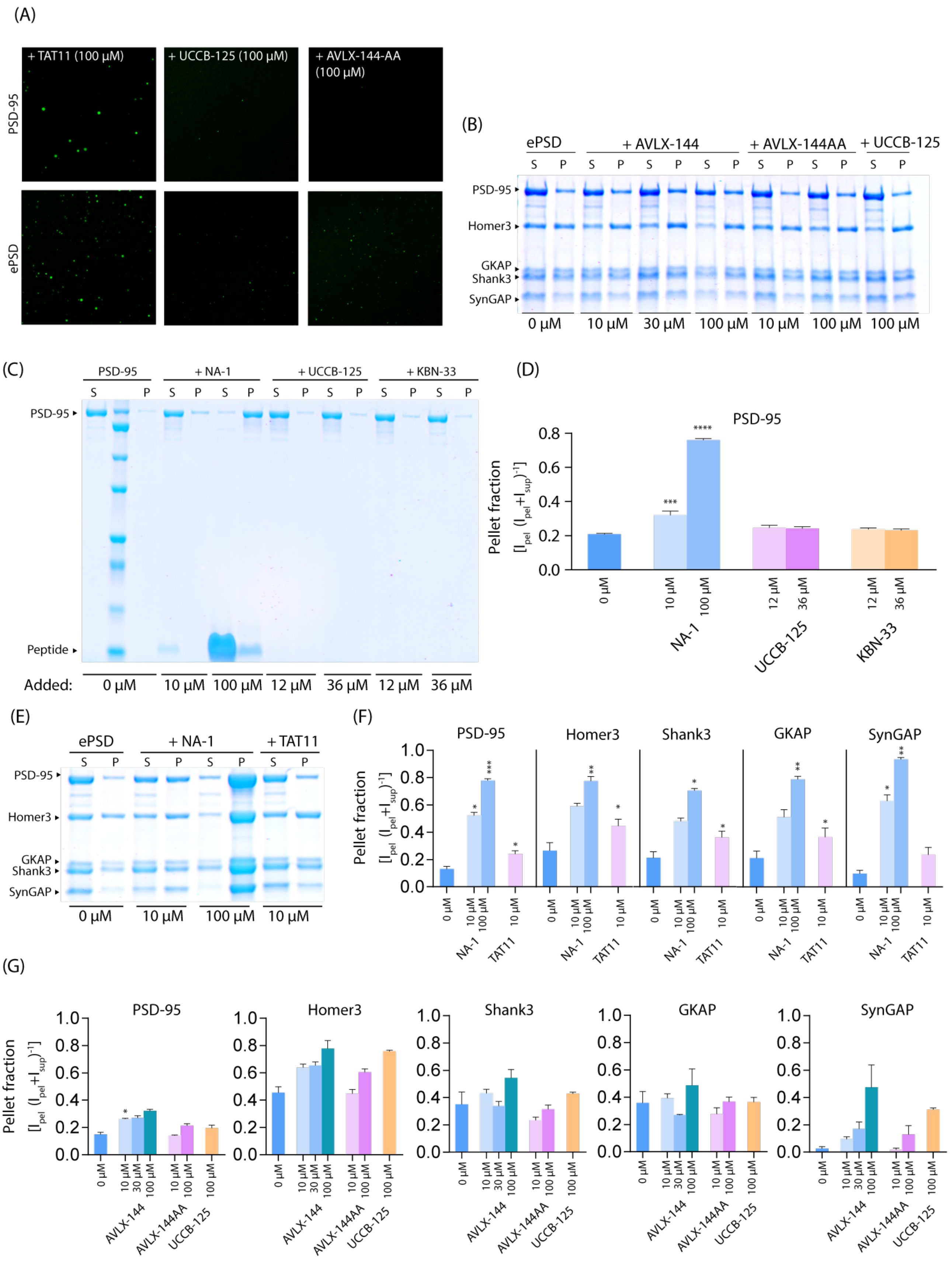
(A)
+ TAT11 (100 µM)
+ UCCB-125 (100 µM)
+ AVLX-144-AA (100 µM)
PSD-95
ePSD
(B)
ePSD
+ AVLX-144
+ AVLX-144AA
+ UCCB-125
S P S P S P S P S P S P S P
PSD-95
Homer3
GKAP
Shank3
SynGAP
0 µM 10 µM 30 µM 100 µM 10 µM 100 µM 100 µM
(C)
PSD-95
+ NA-1
+ UCCB-125
+ KBN-33
S P S P S P S P S P S P S P
PSD-95
Peptide
Added: 0 µM 10 µM 100 µM 12 µM 36 µM 12 µM 36 µM
(D)
PSD-95
Pellet fraction [I_pel (I_pel+I_sup)^-1]
0 µM
10 µM 100 µM NA-1
12 µM 36 µM UCCB-125
12 µM 36 µM KBN-33
(E)
ePSD
+ NA-1
+ TAT11
S P S P S P S P
PSD-95
Homer3
GKAP
Shank3
SynGAP
0 µM 10 µM 100 µM 10 µM
(F)
PSD-95
Homer3
Shank3
GKAP
SynGAP
Pellet fraction [I_pel (I_pel+I_sup)^-1]
0 µM 10 µM 100 µM 10 µM
NA-1 TAT11
(G)
PSD-95
Homer3
Shank3
GKAP
SynGAP
Pellet fraction [I_pel (I_pel+I_sup)^-1]
0 µM 10 µM 30 µM 100 µM 10 µM 100 µM 100 µM
AVLX-144 AVLX-144AA UCCB-125

**Figure S12** (A) Representative images of LLPS droplets for PSD-95 (top) or ePSD (bottom) with indicated peptide. (B) SDS-PAGE sedimentation assay of ePSD (3 µM) upon addition of increasing amounts of indicated peptide. Quantification is shown in main Figure 5E. (C) Representative SDS-PAGE sedimentation assay of PSD-95 incubated with a selection of known inhibitors. NA-1 (Aarts et al., 2002), UCCB-125 (Bach et al., 2009), KBN33 (Nissen et al., 2015). (D) Quantification of (C), shows NA-1 being able to induce LLPS for PSD-95 alone. (E) SDS-PAGE sedimentation assay and quantification (F) of ePSD incubated with indicated peptides, NA-1 and TAT11, both was able to promote LLPS. (G) Quantification of SDS-PAGE sedimentation assay of ePSD incubated with indicated peptides, AVLX-144, AVLX-144-AA and UCCB-125. AVLX-144, AVLX-144-AA (Bach et al., 2012). Error bars are shown as SEM and n=3.

### Supplementary Tables

#### Supplementary table 1. SAXS data collection table

| | PSD-95 |
|---|---|
| **Sample details** | |
| Uniprot ID | P78352 [Human] |
| Buffer | 50 mM Tris (pH 8.2), 300 mM NaCl, 5 mM EDTA, 1 mM TCEP |
| Protein concentration | 30-210 µM |
| Molecular weight | 81 kDa |
| **SAXS data collection details** | |
| Instrument | P12, Petra III, DESY |
| Data collection | 10 May 2019 |
| Wavelength | 1.24 Å |
| Measured q-range | 0.0025-0.73 $Å^{-1}$ |
| Absolute calibration | Water |
| Exposure time | 10 ms |
| Temperature | 283.35 K |
| **Software** | |
| Indirect Fourier transformation to obtain p(r) | GenApp.Rocks |
| Fitting of data with combined analytical and atomic models | EOM, Ranch, Gajoe |
| Data rebin | Will it rebin / Binfactor 1.02 |
| **Model fitting parameters** | |
| Ensemble optimization method (EOM) | |
| Ranch pool generation | 10.000 models / pool, 15 Harmonics, 10% Symmetric structures |
| Gajoe selection and fitting | 1000 generations, 100 ensembles, max 20 curves pr ensemble, 100 repetitions |
